# A systematic review of *Drosophila* short-term-memory genetics: meta-analysis reveals robust reproducibility

**DOI:** 10.1101/247650

**Authors:** Tayfun Tumkaya, Stanislav Ott, Adam Claridge-Chang

## Abstract

Geneticists use olfactory conditioning in *Drosophila* to identify learning genes; however, little is known about how these genes are integrated into short-term memory (STM) pathways. Here, we investigated the hypothesis that the STM evidence base is weak. We performed systematic review and meta-analysis of the field. Using metrics to quantify variation between discovery articles and follow-up studies, we found that seven genes were both highly replicated, and highly reproducible. However, ~80% of STM genes have never been replicated. While only a few studies investigated interactions, the reviewed genes could account for >1000% memory. This large summed effect size could indicate irreproducibility, many shared pathways, or that current assay protocols lack the specificity needed to identify core plasticity genes. Mechanistic theories of memory will require the convergence of evidence from system, circuit, cellular, molecular, and genetic experiments; systematic data synthesis is an essential tool for integrated neuroscience.

## Introduction

Learning is the process by which external sensory experiences and internal states lead to behavioral adaptation. Learning manifests as physiological changes in the brain, the most prominent of which are alterations to the connections between neurons, known as synaptic plasticity (Takeuchi, Duszkiewicz, and Morris 2014). The plasticity theory of learning draws on findings from numerous experimental systems, including vertebrate models, most notably mouse, and invertebrate models, such as the sea slug *Aplysia californica* and the vinegar fly *Drosophila melanogaster* (Bailey, Kandel, and Harris 2015; Ehmann, Owald, and Kittel 2017; Cognigni, Felsenberg, and Waddell 2017). For decades, the *Drosophila* model has been used to characterize the neurogenetic mechanisms underlying memory formation (Ronald L. Davis 2011). Much of our knowledge of the neurogenetics of learning derives from experiments using Pavlovian odor (olfactory) conditioning (Quinn, Harris, and Benzer 1974). Olfactory conditioning uses the simultaneous presentation of an odor paired with an inherently valued stimulus, usually painful electric shocks or a nutritious sugar meal (T. Tully and Quinn 1985; Tempel et al. 1983). Conditioned animals display altered approach/avoidance responses to subsequent odor presentation. When the post-learning trial is performed within minutes of conditioning, the response is referred to as short-term memory (STM). Starting with rutabaga (rut) and dunce (dnc) in the 1980s, such experiments performed on *Drosophila* mutants have identified a number of STM genes (Heisenberg 2003; Keene and Waddell 2007; Tomchik and Davis 2013).

Many STM genes are predominantly expressed in a single brain structure, but an integrated model of the overall STM signaling architecture is lacking. This absence contrasts starkly with other *Drosophila* gene systems. For example, the *Drosophila* signaling networks underlying both embryonic development (Norbert Perrimon, Pitsouli, and Shilo 2012; St Johnston and Nüsslein-Volhard 1992) and the circadian clock (Hardin 2011) have been characterized in detail. In those cases, our molecular-genetic knowledge has reached such an extent that it informs mathematical models that can recreate key system properties (Segal et al. 2008; Fathallah-Shaykh, Bona, and Kadener 2009). It is unknown why genetics has succeeded in defining those aforementioned systems, while delineating similarly complete plasticity pathways has been hard.

Many hypotheses could be put forward to explain why the genetics of memory formation have not yet reached a level of clarity. One such possibility is a weak evidence base. For instance, circadian-rhythm genetics benefits from very large effect-sizes, while memory phenotypes are smaller (Takahashi, Shimomura, and Kumar 2008), making reliable measurements more difficult. Many scientists believe that there is a reproducibility crisis in biomedical research (Baker 2016), and intense investigations into data reproducibility have ensued (“The Challenges of Replication” 2017; Lithgow, Driscoll, and Phillips 2017), including in the fields of cancer research (Begley and Ellis 2012), drug target identification (Prinz, Schlange, and Asadullah 2011), and human psychology (Open Science Collaboration 2015a; Gilbert et al. 2016; Anderson et al. 2016). In medical research, scientists routinely assess a field’s evidence base with statistical synthesis (Haidich 2010). Such meta-research—although relatively rare in the basic biomedical sciences—is growing in importance (“Meta-Analysis in Basic Biology” 2016; Claridge-Chang and Assam 2016; Yildizoglu et al. 2015).

To test the hypothesis that the development of an integrated model of STM has been frustrated by irreproducibility, we evaluated the evidence base with a synthetic analysis of loss-of-function alleles. Our systematic review identified 32 STM genes, of which 23 were amenable to meta-analysis. We applied three metrics across several types of replication to quantify reproducibility. The findings on replicated genes were consistent, and statistical evidence for publication bias was absent. These findings refute our hypothesis and confirm good reproducibility. However, independent replication was rare: most replication was reported in follow-up studies from the discovery group; and only seven of the 32 genes—just 22%—were replicated independently. These results indicate that the *Drosophila* memory-genetics evidence base is divided: a low replication rate for most genes, with robust data integrity for a selected few.

## Materials and Methods

### Information sources and database search

This review was conducted by searching PubMed and Embase databases on 8^th^ April 2017 with the phrase: *“Drosophila* AND (learning OR memory) AND (olfactory OR olfaction OR T-maze OR “T maze” OR odorant OR odor) NOT review[Publication Type]”. The query returned 648 and 559 publications from Pubmed and Embase, respectively. The results were downloaded as .csv files, and the two sources were merged. Publications that were not research articles or not written in English were excluded; this resulted in the identification of 743 articles (Figure 1).

**Figure 1.**
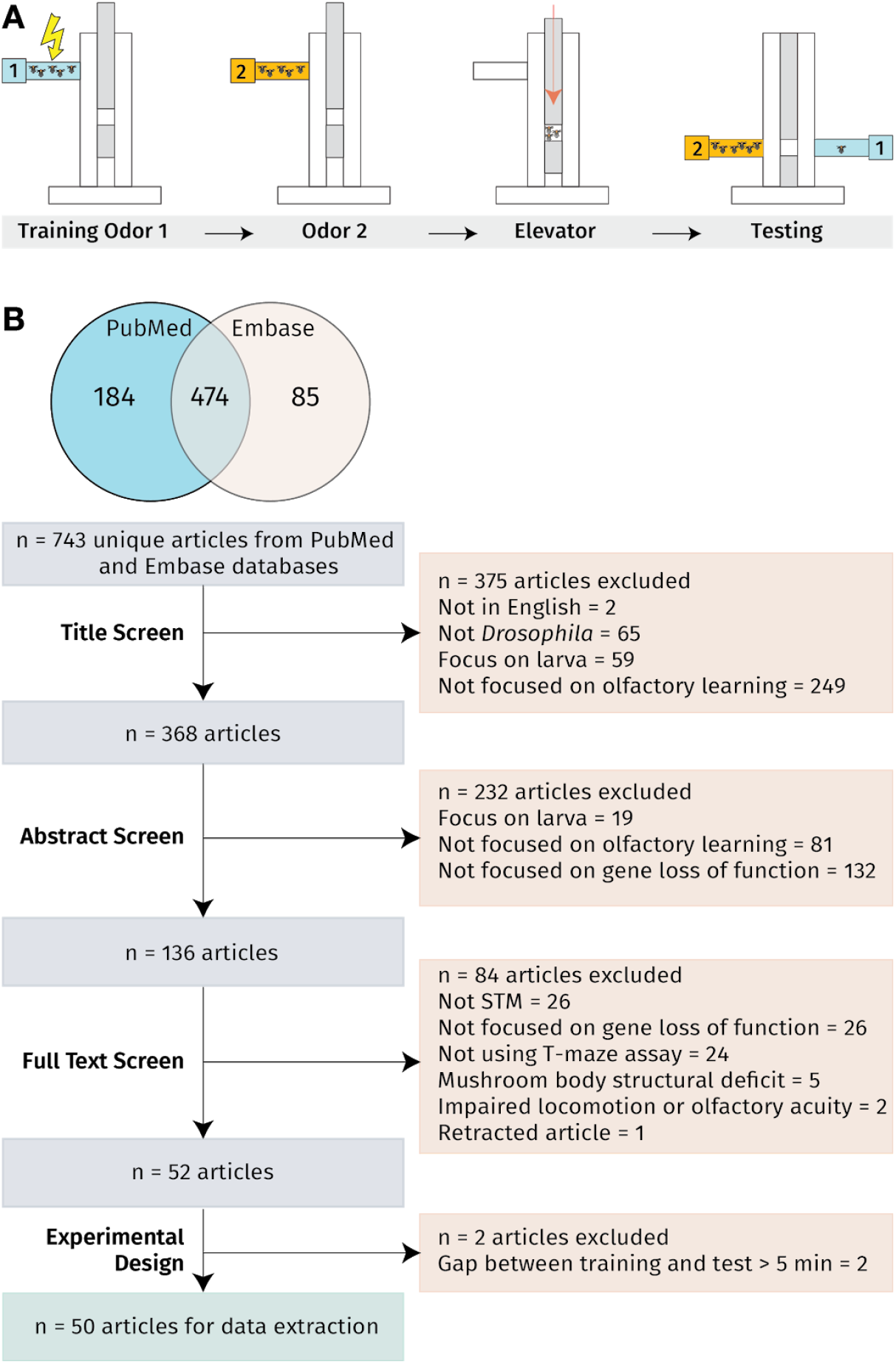
T-maze experiments and the systematic review procedure. **A**. Key steps in the T-maze conditioning protocol. At the start of each conditioning experiment approximately 60-100 flies are tapped into a training tube. Atypical training cycle begins with the presentation of an odor, administered simultaneously with electric shocks. After removal of the shock-paired odor, the flies are presented with a second odor without shock. Subsequently, flies are tapped into the elevator. The elevator with the flies is pushed to the to the choice point position and the two odors are pumped from either end. After two minutes, the elevator is raised to separate the chambers, and the numbers of flies in the chambers are used to calculate a performance index (PI). **B**. PubMed and Embase searches performed in April 2017 identified 743 articles relevant to *Drosophila* short-term memory genes. A four-stage screening process excluded 693 articles, leaving 50 for further review and meta-analyses.

### Eligibility criteria

The systematic review included studies that met the following six criteria: (1) report STM performance defined as ≤ 5 min memory (Heisenberg 2003; Yildizoglu et al. 2015); (2) measure STM in adult *Drosophila melanogaster;* (3) focus on homozygous loss-of-function mutations, including broad RNA interference (RNAi) knockdown mutants; (4) use the T-maze assay; (5) report the full performance index (PI) score for control and experimental flies; and (6) use mutants with intact locomotor activity, olfactory acuity, and brain development processes. The T-maze assay uses electric shock to condition one of two odors, and memory is tested in an odor-choice assay (Figure 1A) (Tim Tully and Quinn 1985). Due to the rapid transition of STM to middle-term memory (Heisenberg 2003), we excluded experiments where the time interval between training and testing was >5 min.

### Study selection

Articles were screened for exclusion in four successive stages: (1) title—375 excluded, (2) abstract—232 excluded, (3) full text—84 excluded, and (4) experimental design—2 excluded. After screening, 50 articles that fully satisfied our selection criteria were taken forward for data extraction and meta-analyses (Figure 1).

### Data extraction

We extracted the following experimental parameters from the 50 included articles: author, year, figure and panel number, experimental genotypes, control genotypes, full PIs of all conditions, standard error of the mean (SEM), sample size (N), temperature, relative humidity, odor pairs, shock voltage, current type, number of training, number of shocks per training, training duration, and the training-testing time interval. If tabulated data were not available, the PI and SEM were extracted from graphs using the Adobe Illustrator (Adobe Systems USA) measurement tool and extrapolated from the y-axis length to obtain a numerical value.

### Summary measures

The PI of the control-group STM can vary considerably between studies (Yildizoglu et al. 2015), which renders inter-study comparisons challenging. As such, we used performance percent change (PPC) as previously described (Yildizoglu et al. 2015). For each experiment, the PPC was calculated as follows:

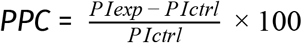

The standard error (SE) of the PPC was calculated using the delta approximation (Cramer 1946; Oehlert 1992; Yildizoglu et al. 2015), as follows:

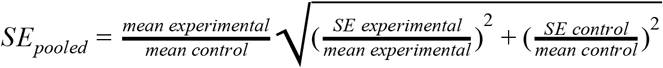

### Outlier exclusion

Outliers were excluded from the dataset based on Z-score values, following best practice (Pagano, Gauvreau, and Pagano 2000; Altman 1968). Effect sizes with a Z-score >2 were removed from the dataset. In total, eight experiments conducted on five alleles were excluded and are greyed-out in the corresponding forest plots.

### Meta-analysis calculations

Where multiple experiments were available (for 23/32 genes), data were meta-analyzed with a random effects model (RE model) to calculate summary effect sizes. Subgroup analyses were also performed on the genes with data for multiple alleles. (When a subgroup had only one experiment, the fixed effects model was used by default.) Both analyses used the metafor package in R (Viechtbauer and Others 2010).

### Publication bias

Some research literatures show a bias in favor of the publication of large effect sizes, and the censorship of small effect sizes (Egger, Smith, and Phillips 1997). Publication bias was examined using funnel plots, which examine symmetry of study effect-size variation around a meta-analytic effect size: asymmetry can indicate bias.

For genes where ≥10 internal replicates were available, publication bias was assessed by constructing a funnel plot of effect sizes (Sterne et al. 2011). Effect sizes and corresponding precisions were plotted against each other and inspected for symmetrical distribution around the meta-analytic mean (Light and Pillemer 1984; Liu 2011). Egger’s method was then applied to test for bias in each gene’s data (Egger, Smith, and Phillips 1997).

### Replication terminology

An *ad hoc* literature search failed to identify definitions of ‘replication’ or ‘reproducibility’ that are broadly accepted and/or statistically formalized. In the context of genetic experiments for STM, we found that *replication* could have at least four meanings: (1) the sample size of a single experiment; (2) multiple experiments reported in a single discovery study; (3) replication reported in a follow-up study by the discovery laboratory; and (4) genuinely independent replication conducted by at least one group of scientists not involved in the original study. By this taxonomy, definitions 1, 2, and 3 represent *dependent replication* and definition 4 represents *independent replication,* which is the most valuable. In the context of loss-of-function genetics, there is the possibility of quantitative and qualitative differences between various alleles; we refer to experiments conducted with an identical allelic state as *allelic replication.* There is also a delineation between efforts to exactly recreate all conditions of the discovery experiment—*direct replication—*and experiments that vary conditions, with the ability to generalize discovery findings—*conceptual replication* (Ioannidis 2012; Makel, Plucker, and Hegarty 2012).

### Reproducibility measures

Published definitions of *reproducibility* vary widely (Patil, Peng, and Leek 2016; McNaught and Wilkinson 1997; Open Science Collaboration 2015a). For the purpose of this meta-analysis, we defined reproducibility as the quantifiable extent of agreement between replicate effect sizes (Open Science Collaboration 2015b). To estimate this, we adopted three statistical methods. Firstly, we employed heterogeneity (I^2^), which measures the proportion of variance that cannot be attributed to sampling error. This measure is widely used in meta-analyses and is precisely defined (Higgins et al. 2003); close agreement between studies produces a favorably low heterogeneity. However, poor precision in the constituent studies of a meta-analysis can also result in low heterogeneity, giving the false impression of good reproducibility. To address this limitation, we also used the mean absolute difference (MAD) between all replicates in a set (see below). Although MAD does not incorporate meta-analytic weighting, it has the benefit of being an intuitive, direct measure of overall discrepancies between means and is not confounded by imprecision. Finally, we generated violin plots to compare discovery effect sizes with replicate effect sizes (Open Science Collaboration 2015b).

### Heterogeneity assessment of reproducibility

Meta-analysts source data from several studies to estimate the summary effect size of an intervention. This procedure also inspects the assumption that the included effect sizes are drawn from a common population. Heterogeneity (I^2^) describes the proportion of meta-analytic variance that is attributable to samples being drawn from different populations, and was calculated using the following formula:

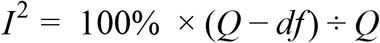

where Q is Cochran’s heterogeneity statistic and *df* is the degrees of freedom (Higgins et al. 2003).

An I^2^ <50% is considered *low*, an I^2^ between 50–75% is considered *moderate,* and an I^2^ >75% is considered *high* heterogeneity (Higgins et al. 2003). Here, we calculated I^2^ for three groupings, namely: 1) all variants of a gene; 2) inter-allelic heterogeneity between the allelic subgroups; and 3) intra-allelic heterogeneity. We used the metafor library in R (Viechtbauer and Others 2010).

### MAD assessment of reproducibility

Discrepancies between all replicates in the units of the effect size can be reported as MAD (also known as Gini’s mean difference) between all effect sizes in a meta-analysis (David 1968). We calculated MAD as follows:

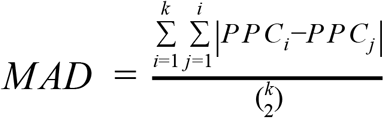

where PPC is the performance percent change, and *k* is the total number of PPCs.

A histogram, QQ-plot, and Shapiro-Wilk test showed that the distribution of effect sizes from the largest meta-analysis (*rut*, Figure 4) were not normally distributed (Shapiro-Wilk P = 0.0015; plots not shown). As MAD is preferred over standard deviation when the data are not normally distributed (Yitzhaki 2002), we adopted the MAD assessment for all gene analyses. Like with I^2^, we used MAD to compare intra-allelic replicates, inter-allelic replicates, and all loss-of-function data for a given gene. Unlike I^2^, there are no pre-established guidelines that relate arbitrary metric thresholds to adjectival descriptors. To place MAD values in context with PPC effect sizes, we chose ≥20% as an arbitrary threshold of *high MAD,* because only 3/23 meta-analyzed genes had effect sizes ≤ |20%| (Figure 3).

### Study protocol and data availability

A study protocol was not pre-registered. Some of the methods were decided upon *a priori,* others were developed during the analysis; this timing is described in a study protocol document available at the Zenodo repository (https://doi.org/10.5281/zenodo.1307126). Data files and code for this study are available from the same repository entry.

## Results

### Experimental conditions vary across studies

We found extensive methodological differences between the 50 published studies (Figure 2). The most commonly used odor pair of 3-octanol, 4-methylcyclohexanol (OCT/MCH) (T. Tully and Quinn 1985) was used to condition flies in <50% of conditioning experiments, while much of the remainder used benzaldehyde combinations. The experimental temperature and humidity settings also varied widely. Most protocols delivered 12 foot shocks (at either 60 V or 90 V); most training periods lasted for 60 s; the shock type (AC or DC) was not reported for half of the protocols (143 of the 278).

**Figure 2.**
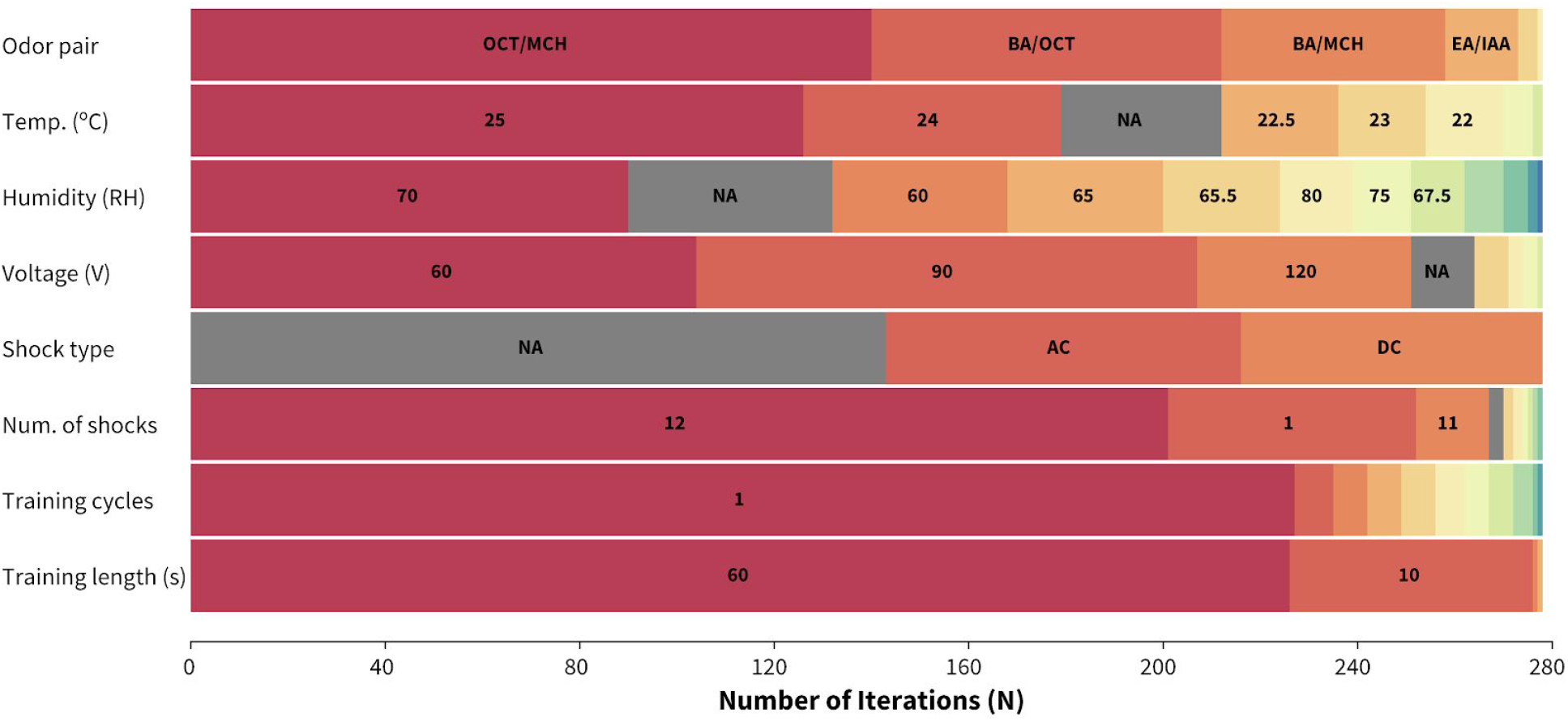
Methodological variation across conditioning experiments. Distribution of experimental conditions across the 278 analysed experiments from 50 articles. EB = ethyl butyrate, EA = ethyl acetate, AA = amyl acetate, IAA = isoamyl acetate, BA = benzaldehyde, NA = not available.

### Most STM studies have not been replicated independently

The reviewed articles implicate 32 genes in *Drosophila* STM; we summarized the meta-analytic results for each gene in Figure 3. Out of the 32 genes, findings on 17 were replicated in at least one follow-up study. However, 10 of these were internal replications conducted by the discovery group; independent replication studies have been conducted for just seven genes. Moreover, only two genes were characterized by >2 independent replicate studies: *rut* (6 replicates) and *dnc* (3 replicates). The other five genes (*Neurofibromin* 1 (*NF*1), rugose (*rg*), Dopamine transporter (DAT), *dopamine 1-like receptor (Dop1R1)* and *fragile X mental retardation* 1 (*Fmr1*)) were replicated by a single independent study. Detailed forest plots and subgroup analyses for individual genes and alleles are shown in Supplementary Figures S1–S15.

**Figure 3.**
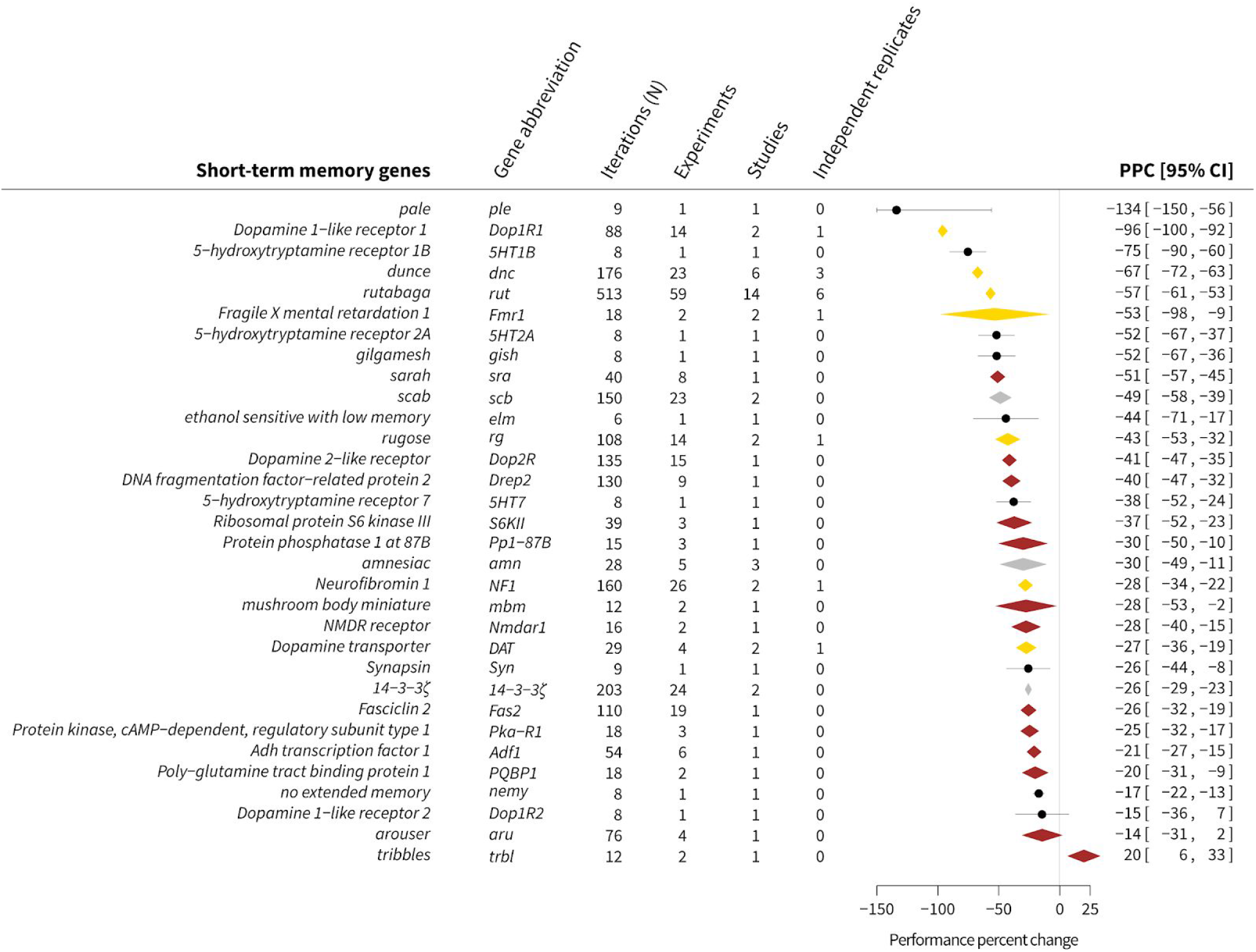
Meta-analytic findings for 32 STM-relevant genes. Summary effect sizes were calculated as performance percent change (PPC) relative to controls (see Methods). Effect sizes reported by independent replications, non-independent replications, and single studies are represented as gold, silver, and bronze diamonds, respectively. Effect sizes reported by one experiment are represented as black dots. Memory alterations in *Drosophila* mutants ranged from +20% (an STM improvement in *trbl* mutants), to –134% (an STM defect in *ple* mutants). The studies show a wide range of iterations, from 6–153. Each iteration reports a full PI score that is derived from ~100 conditioned flies. The total number of experiments for each gene ranges from 1 to 59, where *5HT1B, 5HT2,5HT7, ple, elm, nemy* and *Syn* are represented by only one experiment and *rut* is represented by 59 experiments.

### *Rut* function determines ~57% of STM

*Rut* was one of the first genes implicated in STM (Livingstone, Sziber, and Quinn 1984): it encodes an adenylyl cyclase that generates cyclic adenosine monophosphate (cAMP)(Levin et al. 1992). Extending an earlier meta-analysis (Yildizoglu et al. 2015), our review identified *rut* experiments on 12 loss-of-function alleles and heteroallelic combinations. The STM phenotypes of two alleles (*rut^2080^* and *rut^1^*) have been studied by several groups, but other alleles have not been replicated independently. Two of the *rut* loss-of-function alleles (*rut^769^* and *rut^1951^)* showed a much smaller STM impairment compared to the others (Figure 4). These changes in effect size may be due to different degrees of *rut* deficiency. The aggregate STM reduction caused by loss-of-function *rut* mutations is –57% [95CI –61, –53] with a moderate overall heterogeneity (I^2^) of 65%.

**Figure 4.**
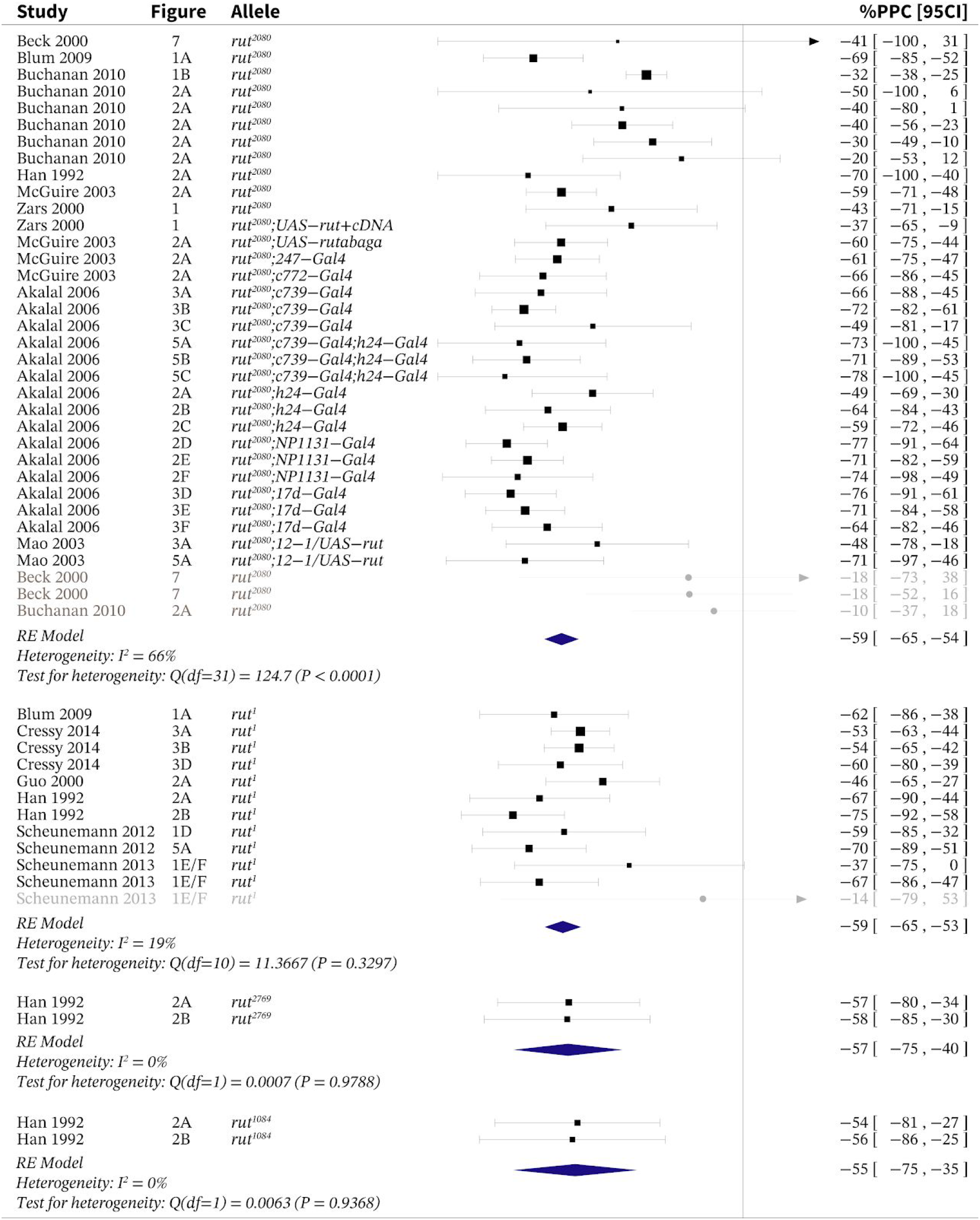
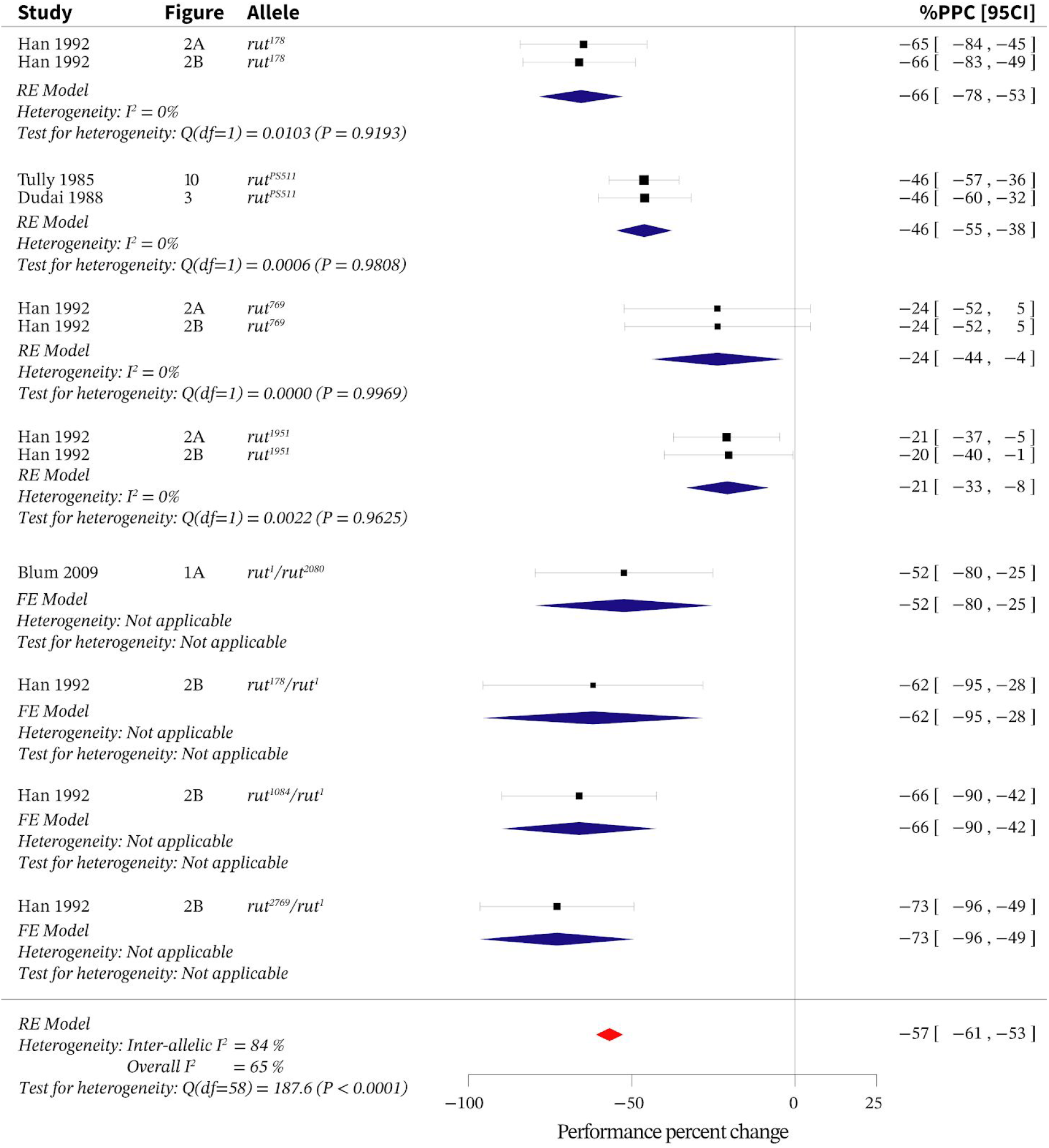
Loss of rut function across different alleles reduces short-term memory (STM) by 57%. Meta-analysis of data from *rut* loss-of-function experiments reveals an overall STM reduction of –57% [95CI −61, −53] with an I^2^ of 65%. Black squares represent the mean performance percent change (PPC) for corresponding experiments; square sizes represent the relative contribution of each experiment to the meta-analytic average. Data sources are indicated in the *study* and *figure* columns; alleles are indicated in their own column. Blue diamonds represent the effect size of the allelic subgroups and the red diamond indicates the overall effect size for *rut.* All error bars (including diamond vertices) represent the 95% CI. The grey-coloured rows indicate the outliers that were excluded from the calculations based on a Z-score outlier filter (see Methods). This data presentation format is repeated for all other forest plots. FE = fixed effects; RE = random effects.

### *Dnc* lesions reduce STM by two-thirds

*Dnc* was the first *Drosophila* gene discovered to modulate memory (Dudai et al. 1976). Subsequent studies have found that *dnc* encodes a cAMP-specific phosphodiesterase that likely consumes the cAMP generated by RUT activity *(R. L. Davis and Kiger 1981).* Here, we identified experiments on seven *dnc* alleles and heteroallelic combinations. The *dnc* meta-analysis revealed a summary effect size of –67% [95CI –72, –63], with moderate heterogeneity, I^2^ = 61% (Figure 5).

**Figure 5.**
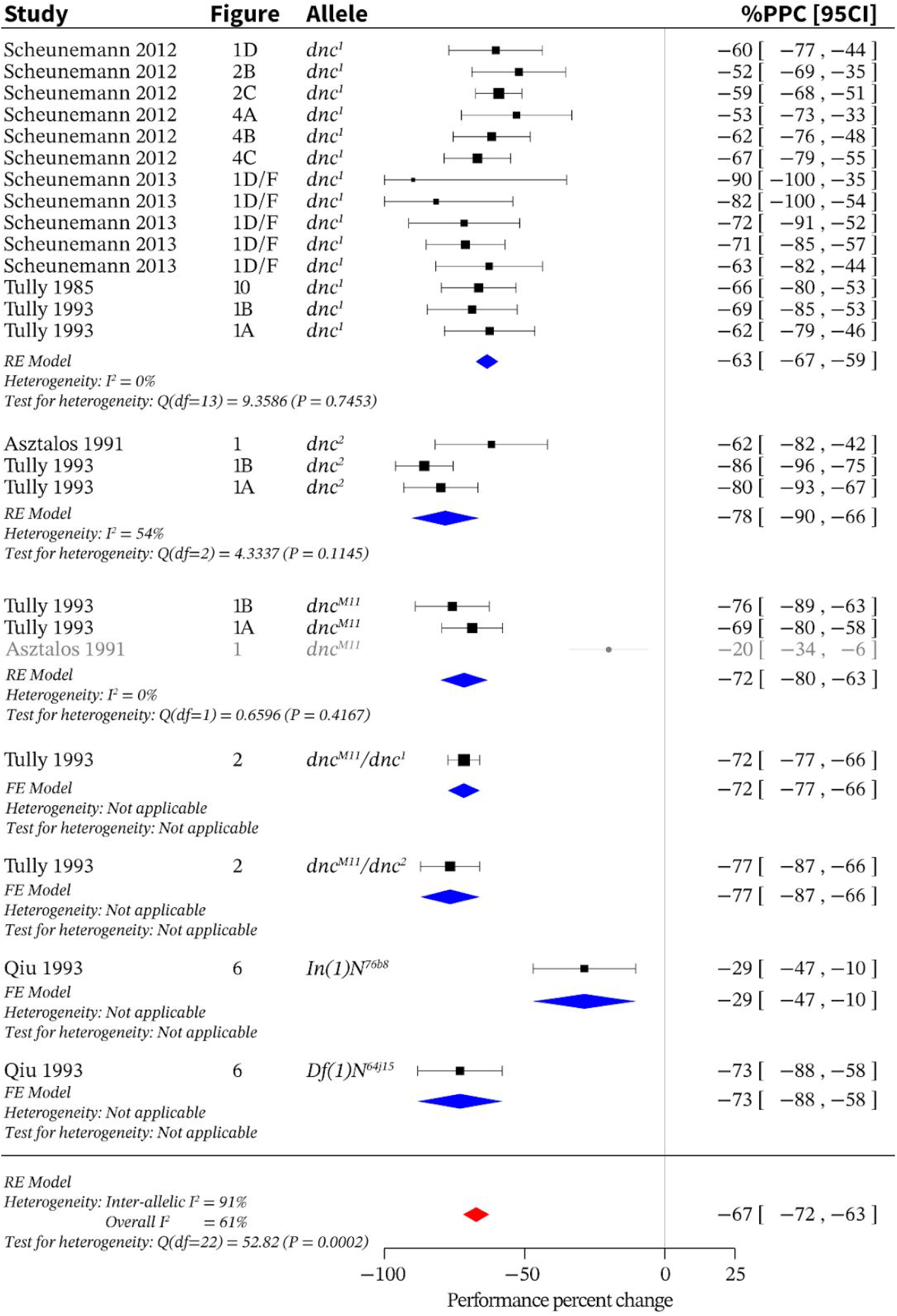
Loss of *dnc* function across different alleles reduces STM by 67%. Meta-analysis of *dnc* loss-of-function data indicates an overall effect size of–67% [95CI −72, −63] with an I^2^ of 61%. FE = fixed effects; RE = random effects.

### *Nf1* function determines ~28% STM

*Nf1* encodes a Ras-specific GTPase activating protein; in humans, mice and flies, *Nf1* lesions elicit various effects including memory deficits (Guo et al. 2000). *Drosophila* NF1 interacts with RUT (Guo et al. 2000) in the mushroom body (MB) (Buchanan and Davis 2010). Here, we identified two STM studies on three loss-of-function *Nf1* alleles: studies on two alleles (*Nf1*^P1^ and *Nf1*^P2^) have been independently replicated but the effect of *Nfl^c00617^* has been investigated only once. Our meta-analysis of all three alleles showed an overall STM decrease of –28% [95CI −34, −22] (Figure 6). However, compared to *Nff1*^p1^ and *Nf1*^p2^ alleles (–31% and –36% respectively), the effect size of *Nf1^c00617^* (–12%) was low. Because the reduction in male body size associated with *NF1* deficiency was also relatively mild in *Nf1*^c00617^ mutants, (Buchanan and Davis 2010), these findings suggest that *Nf1*^c00617^ is a weak hypomorph. As such, the best estimate for the contribution of *Nf1* to STM is about one-third.

**Figure 6.**
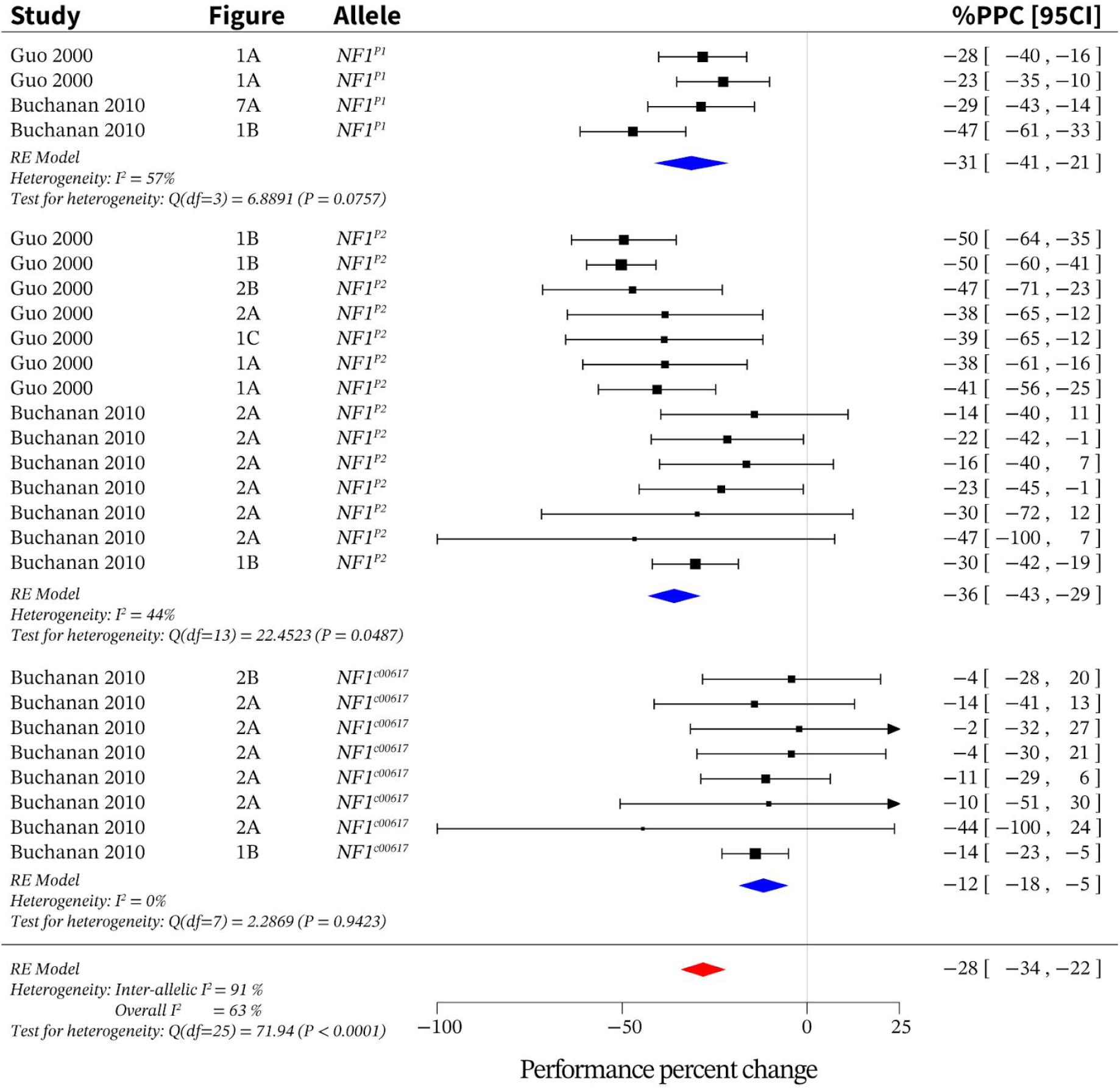
Loss of *Nf1* function across different alleles reduces short-term memory by ~28%. Meta-analysis *of Nf1* loss-of-function mutants produced an overall effect size of –28% [95CI −34, −22] with an I^2^ of 63%. RE = random effects.

### Loss of *rg* function affects STM

*Rg* encodes an A-kinase anchoring protein (AKAP) that is involved in nervous system development (Shamloula et al. 2002). Loss of *rg* function decreases STM, while leaving other types of memory intact (Volders et al. 2012). We identified STM studies on six allelic states: three *rg* hypomorphs (*rg^1^, rg^KG02343^, rg^Y5^),* an RNAi *rg* knockdown, a heteroallelic mutant (*rg^1^, rg^KG02343^)* and one amorph (*rg^FDD^*). The overall decrease in STM was –43% (Figure 7), but only findings on the *rg^1^* allele were independently replicated (Volders et al. 2012; Zhao et al. 2013). Inducing aberrant MB morphology (Volders et a1. 2012; Zhao et al. 2013), the *rg* null has the strongest effect of all *rg* alleles on STM (–89%).

**Figure 7.**
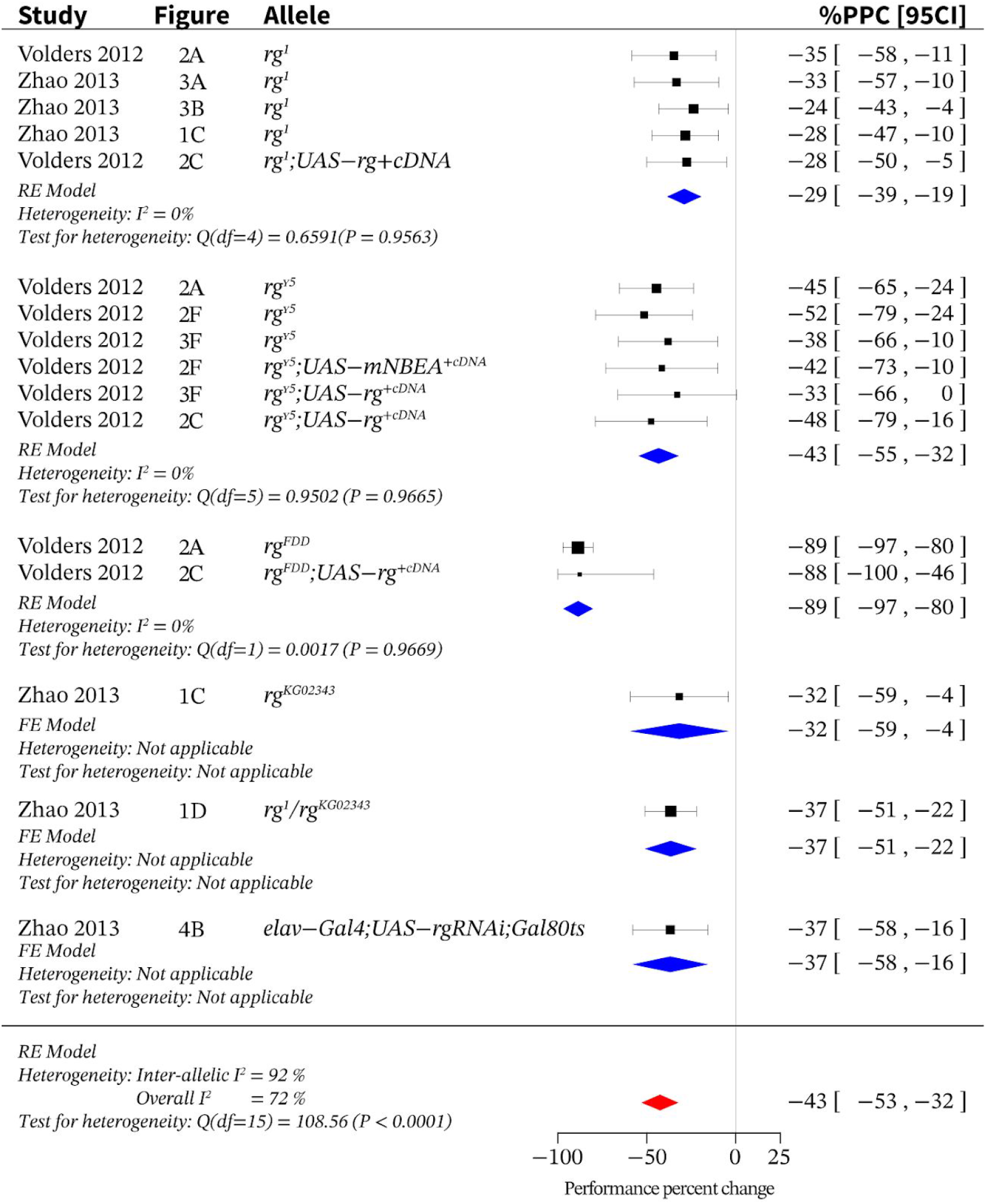
Loss of *rg* function across different alleles leads to an STM reduction of 43%. Meta-ana lysis of *rg* produced an overall effect size of –43% [95CI −53, −32]. Compared to the other alleles, the effect size of the amorphic *rg^FDD^* was more than twice as high: –89%. Heterogeneity across experiments was moderate I^2^ = 72%. FE = fixed effects; PPC = performance percent change; RE = random effects.

### Loss of *Fmr1* elicits a relatively mild STM phenotype

*Fmr1* encodes the fly ortholog of the human FMRP RNA-binding protein associated with Fragile X mental retardation (Y. Q. Zhang et al. 2001). Patients with Fragile X syndrome contain a GGG-triplet expansion in the 5’ untranslated region of *Fmr1,* which causes hypermethylation and transcriptional silencing (Coffee et al. 2012; Verkerk et al. 1991). Our review identified two studies relating *Fmr1* function to STM: one probed the effect of a loss-of-function allele *Fmr1^Δ50M^*, while the other used RNAi to knock down *Fmr*1 (Kanellopoulos et al. 2012; Coffee et al. 2012). The *Fmr1^Δ50^* allele in the homozygous state decreased STM by –77%, while pan-neuronal RNAi knockdown reduced STM by –31% (Figure 8).

**Figure 8.**
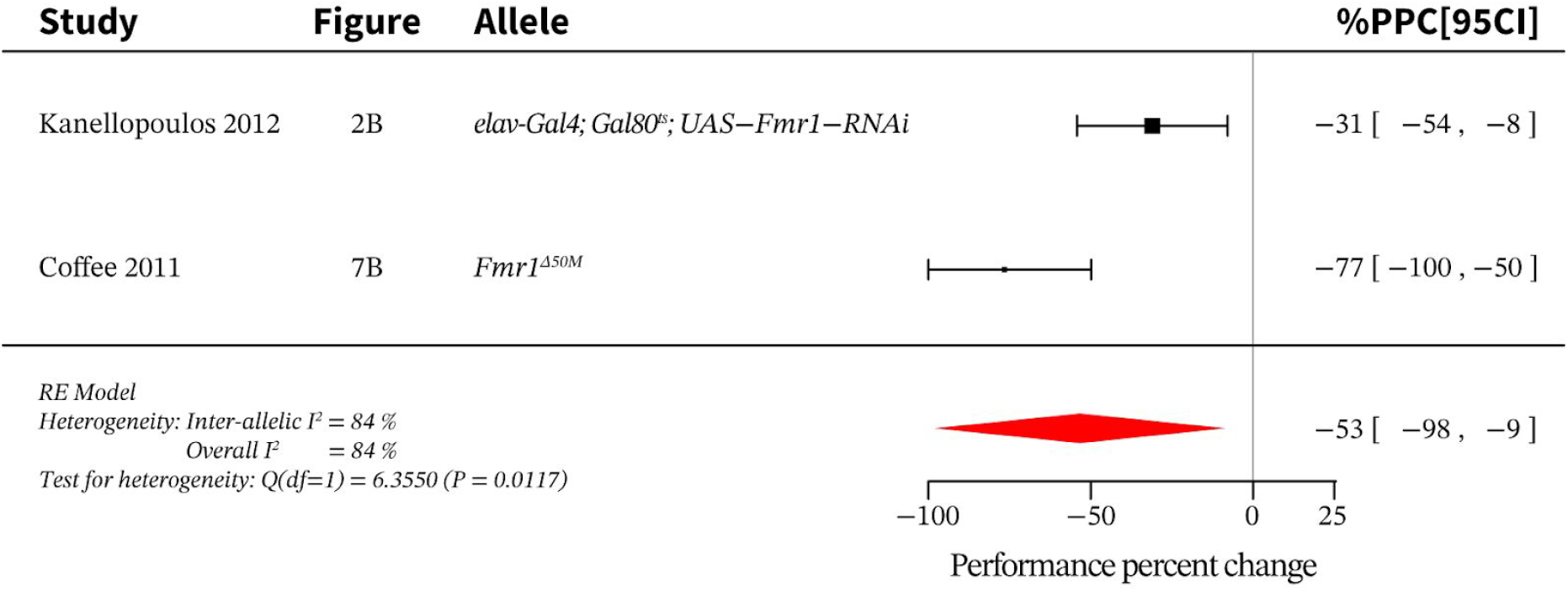
Loss of *Fmr1* across different alleles reduces STM by 53%. Meta-analysis of *Fmr*1 indicates an effect size of –53% [95CI −98, −9]. PPC = performance percent change; RE = random effects.

### Four dopamine-signalling genes affect STM

Dopamine is instrumental to a wide range of behaviours, including STM, and >12 *Drosophila* genes are associated with dopamine metabolism or signaling (Yamamoto and Seto 2014). Of these 12 dopamine-related genes, the systematic review identified STM studies performed on four: *DAT, ple, Dop1R1,* and *Dop2R,* discussed below.

#### DAT

Dopamine shuttles across the plasma membrane via the transporter, *DAT* (S. Zhang et al. 2008; Neckameyer and White 1993; Riemensperger et al. 2011). An imbalance in dopamine levels due to altered DAT function is associated with various neurological disorders and addiction in humans (Yang et al. 2007; S.-I. Ueno 2003). We found two studies (S. Zhang et al. 2008; T. Ueno et al. 2012) that showed that loss of DAT function decreases STM by about –27% (Figure 9A).

#### Pie

The *Drosophila pale* locus encodes the biosynthetic enzyme tyrosine hydroxylase, which is critical for dopamine production. While systemic knockout of *ple* is lethal, one allele abolishes *ple* function exclusively in the central nervous system (CNS), thus permitting viability (Riemensperger et al. 2011). Strikingly, absence of dopamine in the CNS inverts STM polarity: instead of avoiding the shock-associated odors, *ple* mutants actively seek them out—exhibiting a –134% reduction in STM (Figure 3).

#### Dop1R1 and Dop2R

Four different dopamine receptors have been described in flies: *Dop1R1, Dop1R2, Dop2R,* and *DopEcR* (Gotzes, Balfanz, and Baumann 1994; Feng et al. 1996; Hearn et al. 2002; Srivastava et al. 2005). As in mammals, the *Drosophila* D1-type receptors Dop1R1 and Dop1R2 increase cAMP levels upon dopamine agonism (Boto et al. 2014; Beaulieu, Espinoza, and Gainetdinov 2015). Dop1R1 is expressed in the fan-shaped body, the ellipsoid body and the MB; this receptor has important roles in sleep, arousal, and memory (T. Ueno et al. 2012; Andretic, van Swinderen, and Greenspan 2005; Lebestky et al. 2009). In the two studies on this gene (Y.-C. Kim, Lee, and Han 2007; Qin et al. 2012), two *Dop1R1* loss-of-function alleles (*Dop1R1^ln(3LR)234^* and *Dop1R1^f02676^*) almost completely eliminated learning, with an average STM reduction of –96% (Figure 9B). Dop2R is highly expressed in the MB and decreases cAMP levels in response to dopamine agonism (Scholz-Kornehl and Schwärzel 2016). We identified one STM study describing a hypomorphic *Dop2R* mutation that reduced STM by –41% (Scholz-Kornehl and Schwärzel 2016) (Figure 9C).

**Figure 9.**
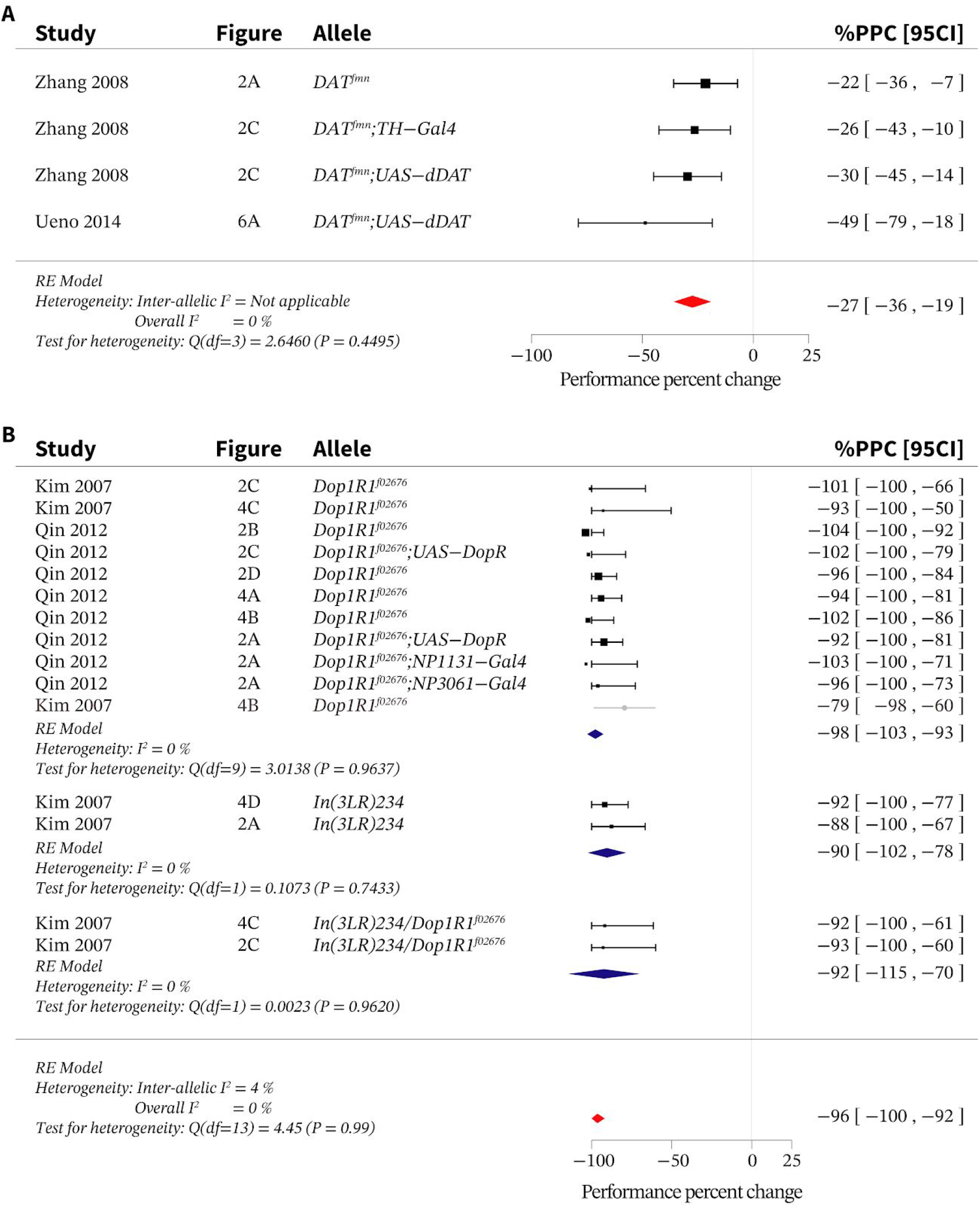
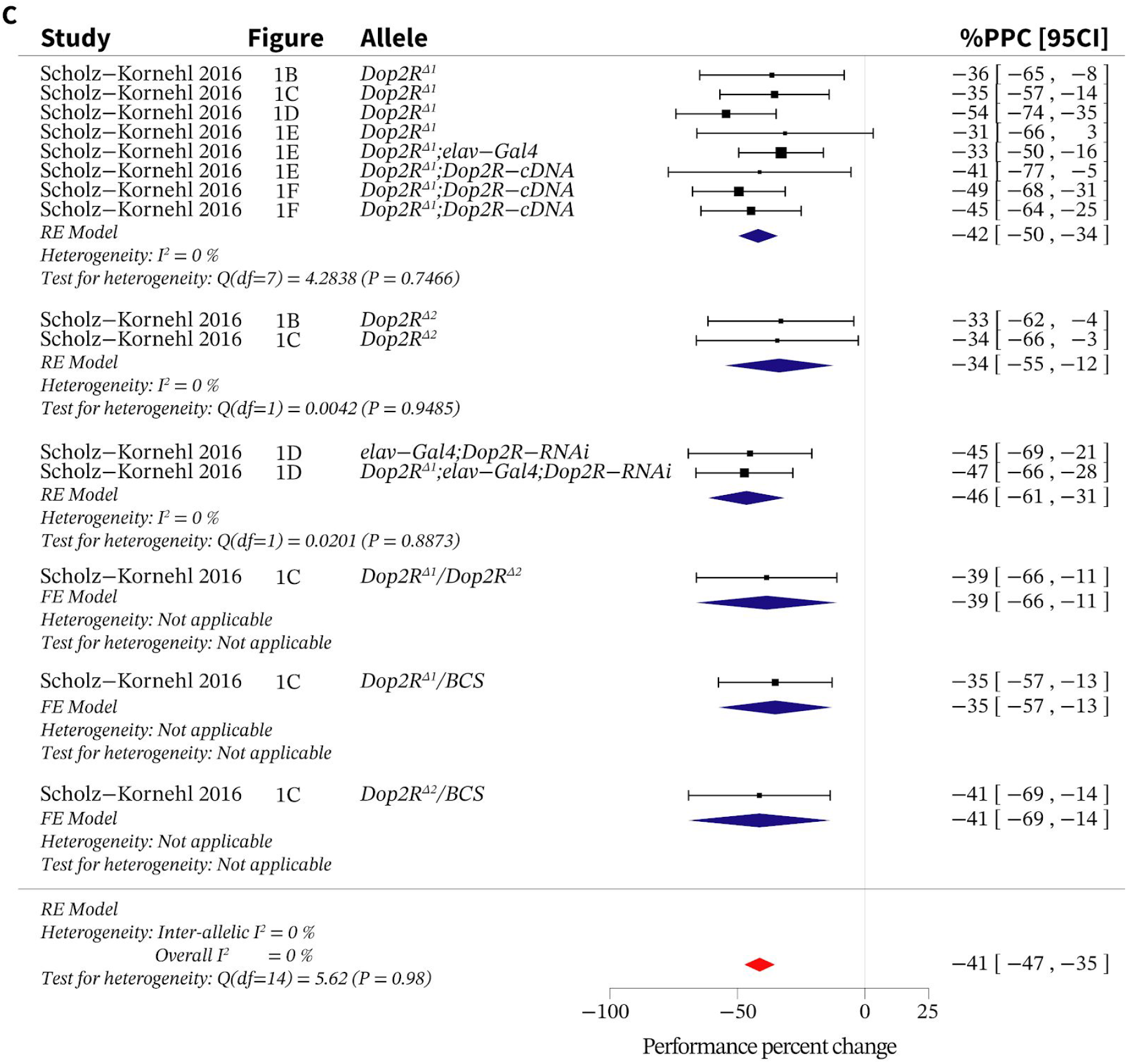
Meta-analyses of genes associated with dopamine signalling. **A**. Meta-analysis of DAT indicates an overall effect size of –27% [95CI −36,-19]. **B**. Meta-analysis of *Dop1R1* indicates an effect size of –96% [95CI −100, −92]. *Dop1R1^In(3LR)234^*, and *Dop1R^f02676^* are also known as *Dop1R1^dumb1^* and *Dop1R1^dumb2^,* respectively. **C**. Disruption of *Dop2R* has an overall effect size of –41%[95CI −47, −35] on short-term memory. PPC = performance percent change; RE = random effects.

### Phosphorylation factors are key to STM

Current hypotheses for the mechanisms underlying memory formation invoke protein phosphorylation and dephosphorylation (Kandel 2012; Margulies, Tully, and Dubnau 2005; Alberini 2009). Five phosphorylation-related genes are associated with STM: *Protein kinase, cAMP-dependent, regulatory subunit type* 1 (*Pka-R1*) (Goodwin et al. 1997); *Ribosomal S6 serine/threonine kinase (S6KII)* (Putz et al. 2004); *Protein phosphatase* 1 *at 87B (Pp1-87B) (Asztalos et al. 1993); Sarah (sra) (Chang, Shi, and Min 2003);* and *gilgamesh (gish) (Tan et al. 2010).* Of these, *Pka-R1* and *S6KII* encode kinases (Kalderon and Rubin 1988; M. Kim et al. 2006), *Pp1-87B* encodes a phosphatase (Baksa et al. 1993), *sra* encodes a calcipressin (a protein that inhibits calcineurin serine/threonine phosphatases) (Chang, Shi, and Min 2003), and *gish* encodes a putative kinase (Hummel et al. 2002). Although no STM discovery studies for these genes have been independently replicated to date, internal replicates permitted meta-analysis of all genes except *gish* (Figures S1–S4; Figure 3). Lesions in these genes produced small-to-moderate STM impairments: *Pka-R1 =* –25%; *Pp1-87B* = –30%; *gish* = –36%; *S6KII* = –37%; and *sra* = –51%. Interestingly, none of these loss-of-function mutations completely abolish STM. Including *Pka-R1*, three of these genes have been characterized in the context of the Rut pathway; *sra* is thought to interact with the cAMP/PKA pathway (Chang, Shi, and Min 2003), while epistasis analysis showed that *gish* STM function is independent of *rut* (Tan et al. 2010).

### STM relies on serotonin receptors

Serotonin (5-hydroxytryptamine; 5-HT) signalling influences numerous *Drosophila* behaviours, including sleep (Yuan, Joiner, and Sehgal 2006; Nichols 2007), courtship (Pooryasin and Fiala 2015), place learning (Sitaraman et al. 2008) and aversive olfactory conditioning. Flies express five serotonin receptors: *5HT1A, 5HT1B, 5HT2A, 5HT2B* and *5HT7* (Johnson, Becnel, and Nichols 2011; Majeed et al. 2016; Gasque et al. 2013). Loss-of-function mutants for three of these receptors *(5HT1B, 5HT2A,* and *5HT7)* have been analyzed by T-maze olfactory conditioning, but no replication studies have been reported. All three mutants exhibited at least moderate learning impairments: a *5HT1B* hypomorph impaired STM by –75%, a *5-HT2A* lesion reduced STM by –52% and a hypomorphic 5-HT7 mutation decreased STM by –38% (Nichols 2007; Johnson, Becnel, and Nichols 2011)(Figure 3).

### STM-relevant genes involved in nucleic-acid function

Three STM genes are associated with nucleic acid function: *Adh transcription factor 1 (Adf1); polyglutamine tract-binding protein* 1 *(PQBP1);* and *mushroom body miniature (mbm). Adf1* is widely expressed during development (DeZazzo et al. 2000), and has a role in synaptic bouton formation in the larval neuromuscular junction (Timmerman et al. 2013). *PQBP1* encodes a polyglutamine tract-binding protein that acts as a transcriptional repressor (Okazawa et al. 2002; Waragai et al. 1999). *Mbm* contains a zinc finger motif—suggesting nucleic acid binding function (Raabe et al. 2004)and is necessary for neuroblast ribosomal biogenesis (Hovhanyan et al. 2014). The effects of lesions in these genes on STM are relatively modest, and have not been independently replicated: *Adf1* = –21%; *PQBP1* = –20%; and *mbm = –33%* (Figures S5–7) (DeZazzo et al. 2000; Tamura et al. 2010; de Belle and Heisenberg 1996).

### Additional intracellular-signaling genes

Five additional intracellular signalling STM genes have been reported: 14*-3-3ζ; Synapsìn (Syn); trìbbles (trbl); arouser (aru);* and *DNA fragmentation factor-related protein 2 (Drep2).* 14-3-3ζ; is preferentially expressed in the MB; loss-of-function alleles produce an STM reduction of –26% STM (Figure S8) (Skoulakis and Davis 1996; Philip, Acevedo, and Skoulakis 2001). *Syn* is a conserved presynaptic phosphoprotein, which among other functions, regulates vesicle recruitment to the readily-releasable pool (Hosaka, Hammer, and Südhof 1999; Rizzoli and Betz 2005); meta-analysis of a study with nine STM loss-of-function experiments showed an STM reduction of –26% (Figure 3) (Godenschwege et al. 2004). A *trbl* lesion uniquely enhanced STM by +20% (Figure S9) (LaFerriere et al. 2008). Hypomorphic *aru* variants have mildly impaired performance during olfactory conditioning, eliciting an STM reduction of only –14% (Figure S10) (LaFerriere et al. 2011). Drep2 is a synaptic protein expressed in the *Drosophila* CNS. Drep 2 expression is especially pronounced at the postsynaptic densities of synapses between projection neurons and Kenyon cells (Andlauer et al. 2014); two *Drep2* deletion alleles decrease olfactory conditioning performance by –40% (Figure S11)(Andlauer et al. 2014). To date, none of the effects of these five genes on STM have been replicated in an independent follow-up study.

### Extracellular-signaling STM genes

Four STM genes were identified with extracellular-signaling functions: *NMDA receptor* 1 *(Nmdar1); Fasci1m 2 (Fas2); scab (scb); and amnesiac (amn).* Of these, only findings on *amn* have been replicated independently. *Nmdar1* encodes a subunit of the NMDA receptor (a heteromeric glutamate-gated cation channel), and is weakly expressed throughout the adult fly brain (Xia et al. 2005). An *Nmdar1* hypomorphic mutation mildly disrupts olfactory learning by –28% (Figure S12)(Xia et al. 2005). *Fas2* is expressed in the MB and is involved in axon guidance and cell adhesion during development (Lin and Goodman 1994; Schuster et al. 1996), and may facilitate dopaminergic input (Cheng et al. 2001). The meta-analysis of 19 experiments from a study on *Fas2* loss-of-function mutants revealed an overall STM impairment of –26% (Figure S13). *scb* is a plasma-membrane α-integrin involved in cell adhesion, and is hypothesized to remodel synapses during learning (Grotewiel et al. 1998). The meta-analysis of *scb* loss-of-function data from 23 experiments found in two studies (Grotewiel et al. 1998; Beck, Schroeder, and Davis 2000) indicated an overall STM reduction of –49% (Figure S14). *Amn* is a putative neuropeptide gene that is expressed in two MB-extrinsic dorsal-paired-medial neurons (Waddell et al. 2000). *Amn* function is required for STM formation; meta-analysis of four experiments in three studies found an overall STM impairment of –30% (DeZazzo et al. 1999; Folkers, Drain, and Quinn 1993; T. Tully and Quinn 1985) (Figure S15).

### STM genes with no known molecular function

Two genes with no-known-molecular function have been implicated in STM: *ethanol sensitive with low memory (elm);* and *no extended memory (nemy). Elm* is predicted to encode a calcium-binding protein, and influences both ethanol sensitivity and STM (LaFerriere et al. 2008); an insertional *elm* allele reduced STM by –44% (Figure 3). *Nemy* is predicted to alter the transcription of neighbouring genes CG8776 and CG8772 ((Kamyshev et al. 2002)); a loss-of-function *nemy* mutation reduced STM by –17% (Kamyshev et al. 2002)(Figure 3). The STM experiments concerning both of these genes have not yet been replicated.

### Replicated findings show no evidence of publication bias

Publication bias is the publication of only positive results and censorship of negative data (Easterbrook et al. 1991). We used funnel-plot analys-s (Egger, Smith, and Phillips 1997) to examine whether the meta-analysed data had a SE-effectsize relationship consistent with publication bias. For this, we selected genes with ≥10 iterations (N) in the meta-analysis (Egger, Smith, and Phillips 1997; Sterne et al. 2011); this selection resulted in nine eligible genes. None of the funnel plots for each gene showed any appreciable asymmetry (Figure 10 A-I). With the caveats of funnel-plot analysis (Sterne et al. 2011), this result indicates that, for the independently replicated STM data, we could find no evidence of bias.

**Figure 10.**
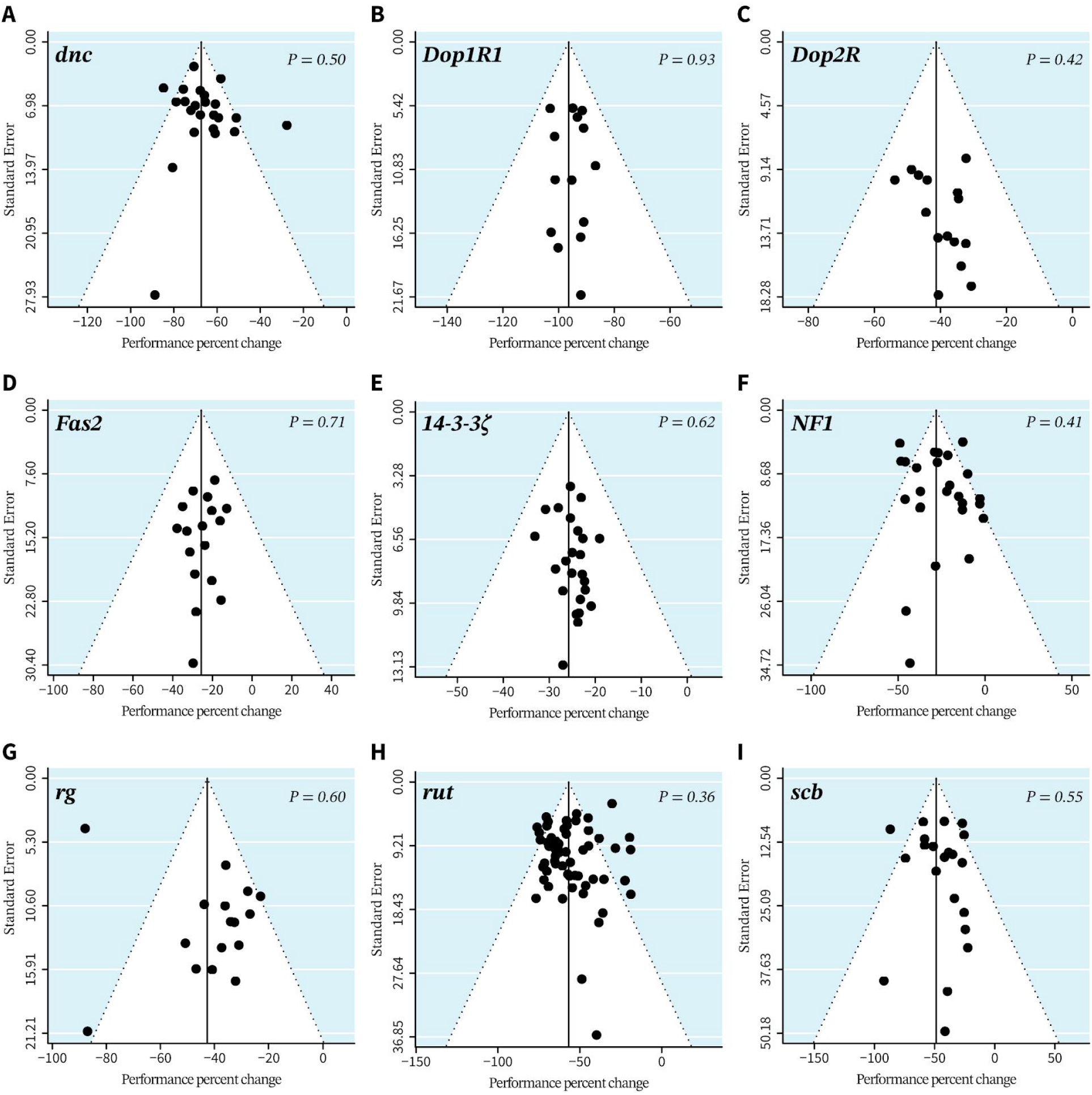
Funnel plot analysis of STM genes shows no publication bias A-I. Comparisons of the standard error (a measure of precision) with the performance percent change (effect sizes from all studies) for nine genes shows largely symmetrical distributions around the meta-analytic summary effect sizes (triangular apex and vertical black line). Egger’s regression tests, which evaluate the significance of the bias (P-values in plot), did not provide evidence of publication bias for all genes plotted.

### Overall heterogeneity across experiments is low

As previously discussed, the degree of variability across experiments in a meta-analysis can be described by the heterogeneity metric (Higgins et al. 2003). Heterogeneity (l^2^%) is closely related to reproducibility—the lower the heterogeneity, the more closely inter-study data agree. We calculated heterogeneity for all STM genes. Considering that combining different alleles may increase heterogeneity, we also conducted—where needed—subgroup analyses of individual alleles or heteroallelic combinations. Of the 23 meta-analysed genes, the majority (17) showed low overall heterogeneity (I^2^ ≤ 50%); only two *(amn* and *Fmr*1) showed high overall heterogeneity (I^2^ > 75%)(Table 1). We observed large differences in inter-allelic heterogeneity among the subgroups of *amn, dnc, Fmr1, NF-1, rg*, and *rut* (Table 1). By contrast, intra-allelic heterogeneity was low or moderate for all genes except for the *aru*^8,128^ allele (I^2^ = 81.36%) (Figure 11A, Supplementary File 2).

**Table 1:** Reproducibility estimates from 23 meta-analyses. Heterogeneity (I^2^) and mean absolute difference (MAD) values calculated for short-term memory-gene meta-analyses. Heterogeneity indicates the proportion of variance not attributable to sampling error alone; MAD indicates the mean difference between replicates. A hyphen indicates the metric is not applicable to the data set.

| Gene | Inter allelic<br>$\%I^2$ | Overall<br>$\%I^2$ | Inter allelic<br>MAD | Overall<br>MAD |
| --- | --- | --- | --- | --- |
| *Adf1<sup>nal</sup>* | 0 | 0 | 27 | 4 |
| *amn* | 88 | 88 | 24 | 24 |
| *aru* | 0 | 50 | 27 | 20 |
| *DAT* | – | 0 | – | 14 |
| *dnc* | 91 | 61 | 50 | 14 |
| *Dop1R1* | 4 | 0 | 3 | 6 |
| *Dop2R* | 0 | 0 | 28 | 8 |
| *Drep2* | 72 | 39 | 39 | 11 |
| *Fas2* | 0 | 0 | 5 | 9 |
| *Fmr1* | 84 | 84 | 46 | 46 |
| *mbm* | – | 0 | – | 1 |
| *Nf1* | 91 | 63 | 1 | 18 |
| *Nmdar1* | 46 | 46 | 13 | 13 |
| *Pka-R1* | 0 | 0 | 5 | 5 |
| *Pp1-87B* | – | 36 | – | 24 |
| *PQBP1* | – | 0 | – | 0 |
| ***rg*** | 92 | 72 | 26 | 19 |
| ***rut*** | 84 | 65 | 37 | 18 |
| ***S6kII*** | 0 | 0 | 48 | 9 |
| ***scb*** | 63 | 46 | 18 | 22 |
| ***sra*** | 35 | 0 | 4 | 15 |
| ***trbl*** | - | 0 | - | 5 |
| **14-3-3ζ** | 0 | 0 | 17 | 3 |

### No relationship between effect size and allelic heterogeneity

Because small changes in STM are harder to detect than large changes, we hypothesized that genes with subtle performance changes would also have poor reproducibility. To test this hypothesis, we plotted the intra-allelic heterogeneities against allelic effect sizes. We found no relationship between effect size and heterogeneity (adjusted-R^2^ = −0.016, P = 0.6169, Figure 11A).

### Increased replication has a minor effect on allelic heterogeneity

As most studies on STM genes have not been replicated, we next asked whether allelic heterogeneity increased when data were derived from independent replications. To test this hypothesis, we grouped the heterogeneity scores according to the allele’s replication status (for classification see Figure 3). As expected, alleles with a higher replication status generated more heterogeneous results compared to those with a lower replication status (Figure 11B). Intra-allelic heterogeneity in datasets from independently replicated genes (median = 9.4) was substantially higher than both the heterogeneity of the shared-authors replicates (median = 0) and the within-study replicates (median = 0).

**Figure 11.**
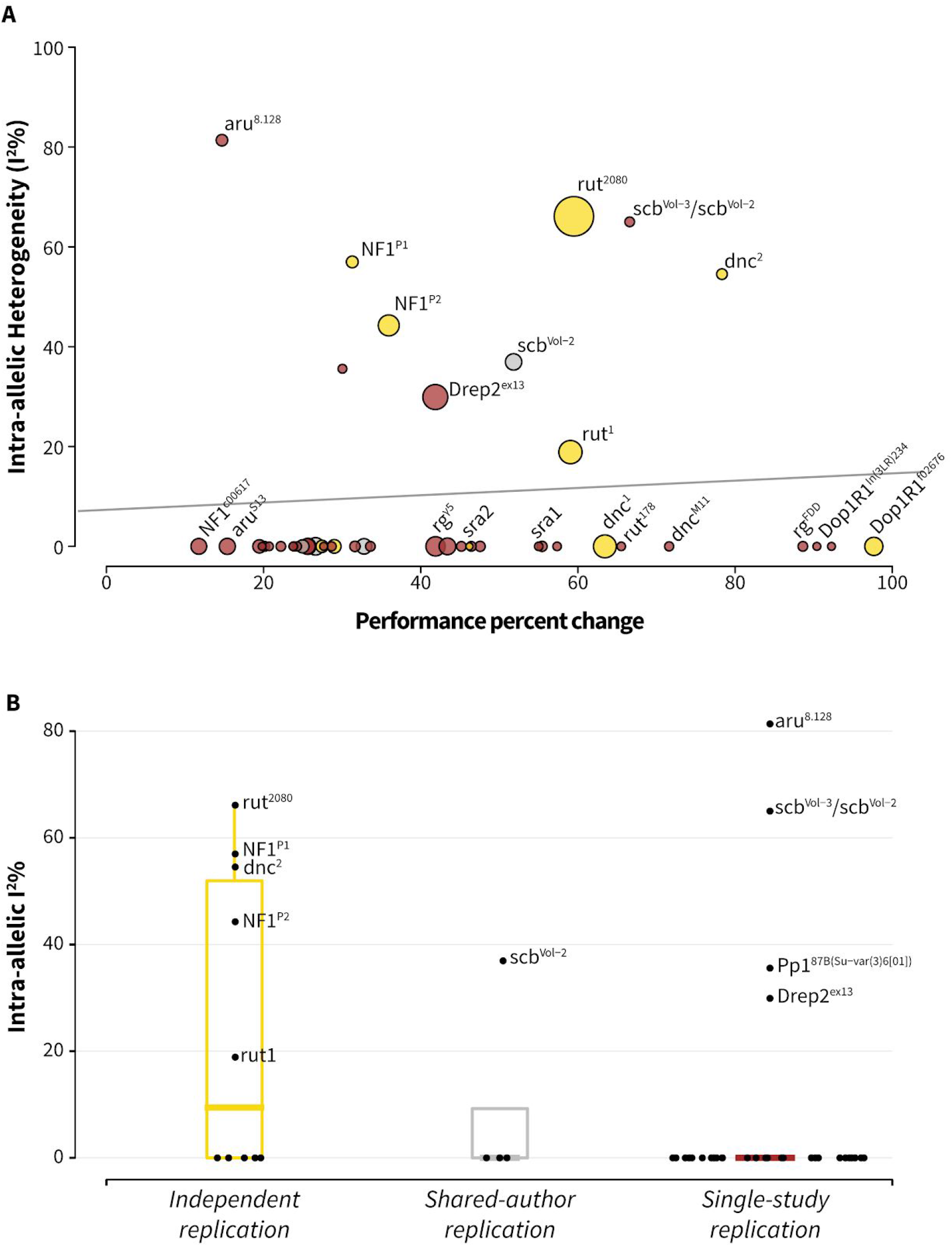
Heterogeneity distribution in the meta-analysed data set. **A**. Effect size does not correlate with intra-allelic heterogeneity (Adjusted-R^2^ = −0.016, *P =* 0.6169). Short-term memory genes are shown as bubbles, with diameters proportional to their sample size and labelled according the replication status (see Figure 3). **B**. Internally replicated results show a lower heterogeneity (I^2^) than those with independent replications. I^2^ medians for independent replicates = 9.441 (interquartile range, IQR 0 to 59.96), shared-authors = 0 (IQR 0 to 9.24), and single-article = 0 (IQR 0 to 0).

### Effect-size reproducibility is high

Due to the drawbacks associated with I^2^ (see Methods), we also calculated the MAD as an independent estimate of reproducibility; MAD is not biased by the SE. For all tiers of replication, the majority of STM genes had a low MAD of <20% (see Methods). Defining high MAD as >20%, only five of the 23 meta-analyzed genes had a high overall MAD: *amn, aru, Fmr1, Pp1-87B,* and *scb* (Table 1). Similarly, for inter-allelic and intra-allelic reproducibility, only ten and five genes had a high MAD, respectively (Table 1, Supplementary File 2). There was no relationship between intra-allelic MAD and the effect size (adjusted-R^2^ = –0.02, P = 0.789), further supporting the conclusion that small effect sizes are not harder to reproduce (Figure 12A). Between the three replication categories, we observed only subtle differences between the median MAD scores (Figure 12B). Both heterogeneity and MAD measures support the idea that *Drosophila* STM studies have good reproducibility.

**Figure 12.**
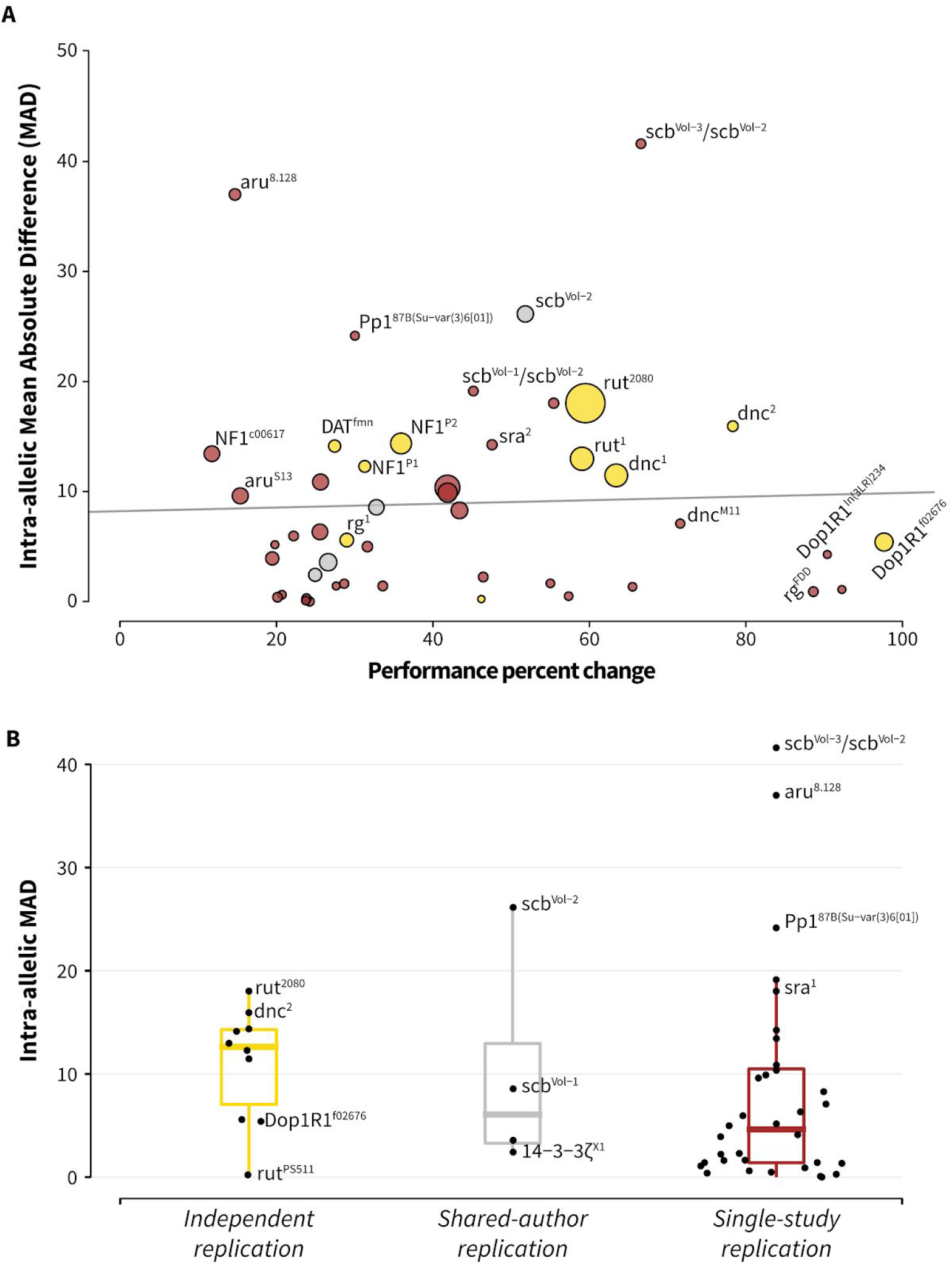
Analysis of reproducibility by mean absolute difference (MAD) scores. **A**. Effect size does not correlate with MAD scores (Adjusted-R^2^ = −0.02058, *P* = 0.789). Short-term memory genes are shown in bubbles that have diameters relative to their sample size, and colors depending on the replication status (gold = independent, silver = shared-author, bronze = single-study.) **B**. Internally replicated results show a somewhat lower MAD score than those replicated independently. Medians of the three tiers were only slightly different, and had substantial overlap: independent = 12.63 (interquartile range, IQR 7.05 to 14.3); same-group = 6.07 (IQR 3.28 to 12.96); and single-article = 4.99 (IQR 1.34 to 10.36). PPC = performance percent change.

### Original findings and independent replications have similar effect sizes

In psychological science, effect sizes decline substantially between discovery studies and follow-up articles (Open Science Collaboration 2015b). For the seven STM genes that have been independently replicated, we compared the average effect sizes from the earliest report for each gene with the available replicate effect sizes. In total, 31 experiments from discovery studies were matched with 105 experiments from follow-up studies. Surprisingly, the later T-maze effect sizes were, on average, +19% higher than those reported in earlier publications (Figure 13). This result refutes the hypothesis of low reproducibility, and supports the idea that STM analyses in *Drosophila* are highly reproducible.

**Figure 13.**
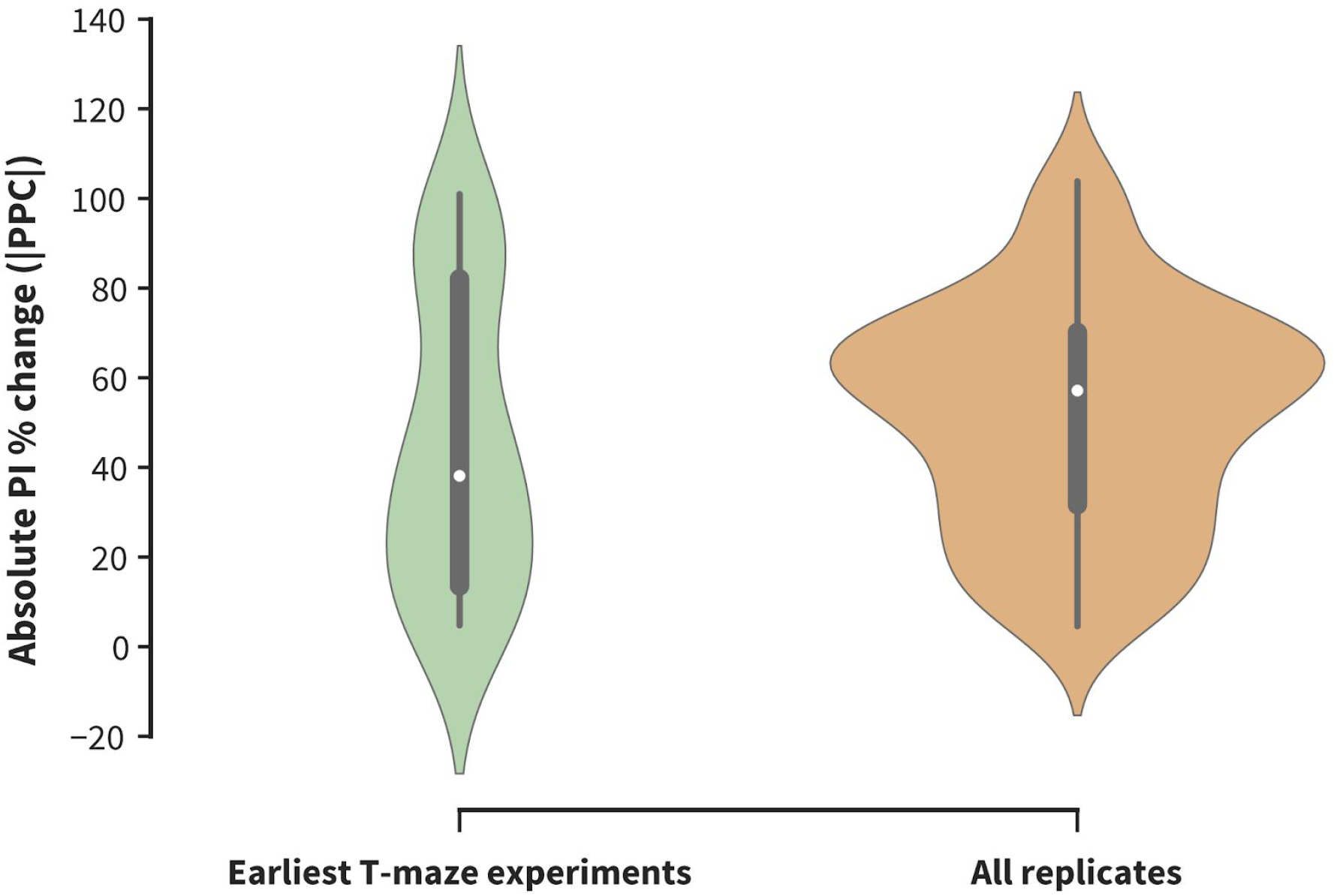
Effect sizes between original and follow-up findings are comparable. Absolute performance percent change (|PPC|) values of the earliest T-maze experiments (green) and their independent replicates (orange) were compared. Experimental data are derived from genes for which replicate experiments are available: *rut, dnc, NF1, Dop1R1, rg, Fmr1, DAT.* The replicate effect sizes are slightly larger than the original effect sizes: median |PPC| for the earliest T-maze studies = 38.12 (interquartile range, IQR 13.54 to 82.07); median |PPC| for all replicates = 57.14 (IQR 31.66 to 70.13), U = 1373.0, *P* = 0.09.

## Discussion

### Meta-research aspects

Recent meta-research debate has focused on the poor reproducibility of data observed in a number of fields, including the psychological sciences (Open Science Collaboration 2015b), and preclinical neurosciences (Button et al. 2013), and how the publication of underpowered experiments is driven by incentive structures (Nuzzo 2015; Munafò et al. 2017; Rosenthal and Rosnow 2009). While widespread irreproducibility is clearly an important problem in many fields, our data support that this does not hold true for *Drosophila* genetic studies of olfactory STM. Although publication bias is found in other areas of neuroscience (Sena et al. 2010; Mohammad et al. 2016), funnel plot analysis indicated that STM gene effect-size distributions were unbiased. Moreover, three quantitative measures of reproducibility (heterogeneity, MAD, and effect-size decline) indicated generally high reproducibility. Thus studies in this field appear more reproducible than other areas of brain science.

There are at least two possible interpretations as to why this finding should be. One positive hypothesis would be that the *Drosophila* STM field has superior data integrity due to the low cost of experiments that allow the use of large sample sizes. Low costs should also mean that independent replication is relatively accessible to other labs, so discovery authors—operating with this knowledge—might favor waiting to publish when key results have been extensively internally replicated. Certainly, our systematic review found ample evidence of extensive internal replication, supporting this positive view. A second, more skeptical, hypothesis points at the relative dearth of independent replication: of the 32 identified genes, only seven have been independently replicated thus far. The bulk of STM loss-of-function replications have been conducted on *dnc* and *rut* mutants, which were both originally identified in the early 1980s (Duerr and Quinn 1982; Livingstone, Sziber, and Quinn 1984). For each non-replicated gene, this view proposes either that there is insufficient interest for any other lab to perform a replication experiment, or that an experiment was conducted but never published. This skeptical perspective holds that the replication shortage could be a cryptic form of publication bias. Indeed, our review found no refutation studies, suggesting that the field abandons—rather than refutes—irreproducible memory genes. This abandonment hypothesis would ideally be tested with pre-registered experiments, as was done in psychology (Open Science Collaboration 2015b).

### Limitations

There are two notable limitations to this study. First, a challenge for all meta-analyses is to balance data splitting and clumping to appropriately account for experimental differences. Direct replication is a noble ideal; in practice however, all follow-up experiments are conceptual replicates to varying degrees as there are always differences between experiments. Here, we addressed experimental variation using sub-groups to examine allelic differences, while combining data from all (non-outlier) loss-of-function alleles for a gene. This approach accommodates different alleles, while yielding a single estimate for the overall impact of each gene. We observed that the literature contains little information regarding the severity of individual alleles, or the qualitative or quantitative differences between alleles; reports of quantitative measures of allelic severity (e.g. q-PCR, ELISA, immunohistochemistry analyses) were the exception (Qiu and Davis 1993; Han et al. 1992; Grotewiel et al. 1998; Goodwin et al. 1997; Chang, Shi, and Min 2003). Due to this absence, multilevel models that might be constructed to account for this variation are inaccessible (Yildizoglu et al. 2015). Despite this limitation, we view the present method as preferable to narrative review, because meta-analyses are systematic and quantitative.

Second, there remains no optimal method, to date, to quantify reproducibility. As significance testing is now deprecated (Gardner and Altman 1986; Halsey et al. 2015; Cumming and Calin-Jageman 2016; Claridge-Chang and Assam 2016), assessing reproducibility with significance test results was avoided (Open Science Collaboration 2015b). Nevertheless, the three methods used here also have limitations: heterogeneity incurs the issues described above; MAD is not standardized, so cannot be compared across experimental systems; and effect-size decline can only provide an overview. We propose that variance-type measures of reproducibility are preferable to significance-test-result methods, which inherently introduce arbitrary threshold distortion (Halsey et al. 2015; Yildizoglu et al. 2015).

### Genetic aspects

Our meta-analyses produced estimates of the quantitative phenotypes for all STM alleles. An advantage of model-system genetics is the ability to assess two gene lesions in combination; this approach allows geneticists to determine whether such combinations are additive or epistatic. Additivity suggests that two genes function independently, while epistasis (sub-additive or super-additive effect sizes) can mean that they operate in the same pathway. The 32 STM genes could be crossed into 1,024 possible two-way combinations; of these, 164 pairs have an effect-size sum exceeding 100%. Despite this, we detected only a few studies that contained experiments on trans-allelic combinations, including *rut-dnc* (Scheunemann et al. 2012); *rut-gish* (Tan et al. 2010); and *rut-NF1* (Guo et al. 2000). Although looking for epistasis experiments was not an original goal of the review, the observation (of a systematic set of articles) suggests that integrating existing genes into pathways is largely missing from the field. Including the oddball effect sizes *(ple* and *trbl*), the sum of absolute effect sizes for STM genes is 1309% (Figure 3). Assuming that most gene effects are reproducible, it is difficult to account for this memory over-abundance. There are at least two possible explanations. First, many genes may fall into the same pathways: for example, there could be three pathways, each accounting for ~33% of STM and comprising roughly a dozen genes. Second, it may be that the current methods of measuring STM lesions lack specificity, and that disrupting numerous neuronal processes have multifarious effects on both memory formation *per se* and its preconditions.

All STM-gene-discovery studies demonstrated that the genetic lesion(s) disrupted memory while leaving odor sensitivity, shock sensitivity, and motor function unaffected (Supplementary File 1). These controls establish that a lesion does not affect the constituent sensory-motor systems (Mihalek, Jones, and Tully 1997), with the aim of establishing a specific role in associative functions. Nevertheless, apart from memory, the identified STM genes are reportedly involved in numerous additional biological processes. For example, loss of *NF1* function shortens lifespan, increases sensitivity to oxidative stress (Tong et al. 2007), disrupts circadian rhythms (Williams et al. 2001), and produces excessive grooming behaviours (King et al. 2016). The classical STM genes rut and *dnc* each have pleiotropic effects on at least five non-memory behaviours (Zhong and Wu 2004; Tong et al. 2007; Venkatesh et al. 2001; Chen and Ganetzky 2012; Donlea, Ramanan, and Shaw 2009; Siegel and Hall 1979; Gailey, Jackson, and Siegel 1984; Hong et al. 2008; N. Perrimon et al. 1986; Kiger and Salz 1985; McBride et al. 1999; Kubli 2003). That all STM genes are broadly pleiotropic raises the question as to whether such factors can be considered memory genes *per se*, or should be viewed as neuronal-function genes, perhaps with selective importance to memory cells.

The classic example of a preconditional memory lesion is one that disrupts normal MB development, but is not acutely involved in the physiological changes that occur during STM formation. In addition to olfactory-avoidance controls and neuroanatomy, this confound can be addressed, in part, with temporal control of gene function/dysfunction (S. E. McGuire, Le, and Davis 2001; Dubnau et al. 2001; Sean E. McGuire et al. 2003). However, even this protocol cannot differentiate between genes that have an immediate role in associative plasticity and those that are acutely essential to normal memory-cell function. If the latter type are preferentially expressed in the memory cells, they would be almost indistinguishable from core plasticity factors.

From the 32 years of 50 STM genetics papers, only one was published in the review’s most recent three years, suggesting declining efforts in this area (Supplementary Figure 16). However, the emerging circuit-analysis field retains a considerable reliance on the classic learning-gene literature (Hige et al. 2015; Cohn, Morantte, and Ruta 2015). Nevertheless, a complete theory of memory will require the integration of evidence from system, circuit, cellular, molecular, and genetic analyses; memory genetics remains a topic of crucial importance. The meta-analysis presented here provides a systematic account of STM genetics, and recasts a traditionally significance-testing field in terms of effect-size estimation; such quantitative perspective will be equally important to circuit studies.

## Conclusion

This study investigated the hypothesis that *Drosophila* memory genetics has limited reproducibility. To address this question, we performed a systematic review of the *Drosophila* STM genetics field, defined a taxonomy of replication types, performed meta-analyses, and applied several reproducibility metrics. The resulting synthesis does not support the hypothesis but instead indicates that replicated STM gene discovery experiments are highly reproducible. We propose that the high reproducibility of *Drosophila* behavior data may derive from the low cost per animal, which permits extensive internal replication. However, despite the low cost and prevalence of extensive (internal) replication, this does not appear to have translated into a high rate of independent replication. The reason for this remains unknown. Total current STM-gene lesions have an estimated sum of deleterious effects of ~1300%. Assuming their effects are all reproducible, this finding suggests either that many of the genes fall into shared pathways, or that current protocols lack specificity to identify core associative-memory plasticity factors.

## Acknowledgments

The authors would like to thank Pryseley Assam (Duke-NUS Medical School) for suggesting mean absolute difference, Malcolm Macleod (University of Edinburgh) and an anonymous reviewer for helpful suggestions, and Insight Editing London for critical review and editing of an earlier version of the manuscript.

## Author contributions

*Conceptualization:* ACC; *Methodology:* TT, ACC; *Software:* TT (R and Python); *Data Analysis:* TT; *Writing - Original Draft:* TT, SO; *Writing - Revision:* TT, SO, ACC; *Visualization:* TT; *Supervision:* ACC; *Project Administration:* ACC; *Funding Acquisition:* ACC.

## Supplementary figures

**S1.**
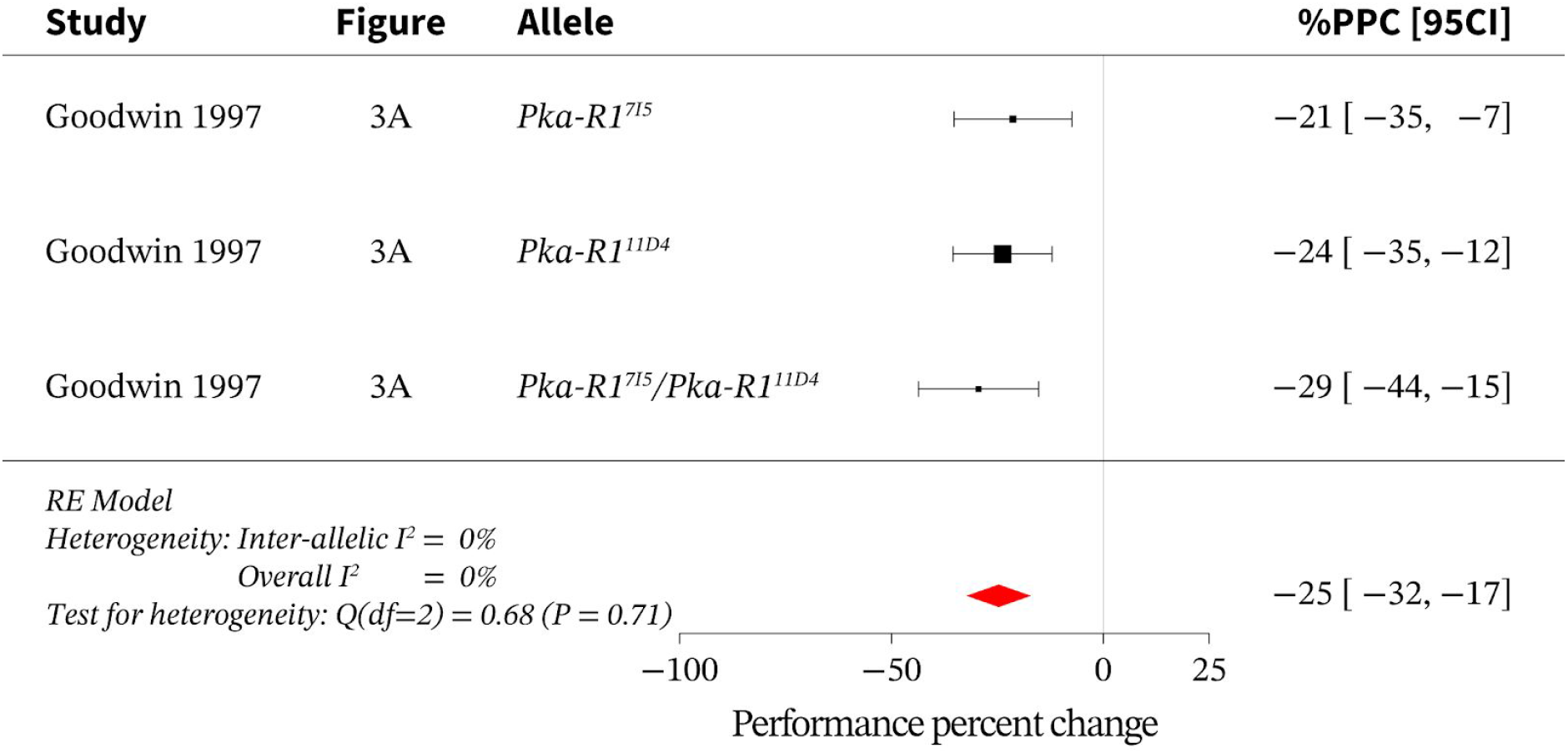
Meta-analysis of loss of function *Pka-R1* alleles. Meta-analysis of *Pka-R1* loss-of-function data indicates an overall effect size of –25% [95CI −32, −17] with an overall I^2^ of 0%. PPC = performance percent change; RE = random effects. Black squares represent the mean performance percent change (PPC) for corresponding experiments; square sizes are relative to each experiment’s weight in the meta-analytic average. Data sources are indicated in the *study* and *figure* columns; alleles are indicated in their own column. The red diamond indicates the overall effect size for *Pka-R1.* All error bars (including diamond vertices) represent the 95% CI. This data presentation format is repeated for all other forest plots.

**S2.**
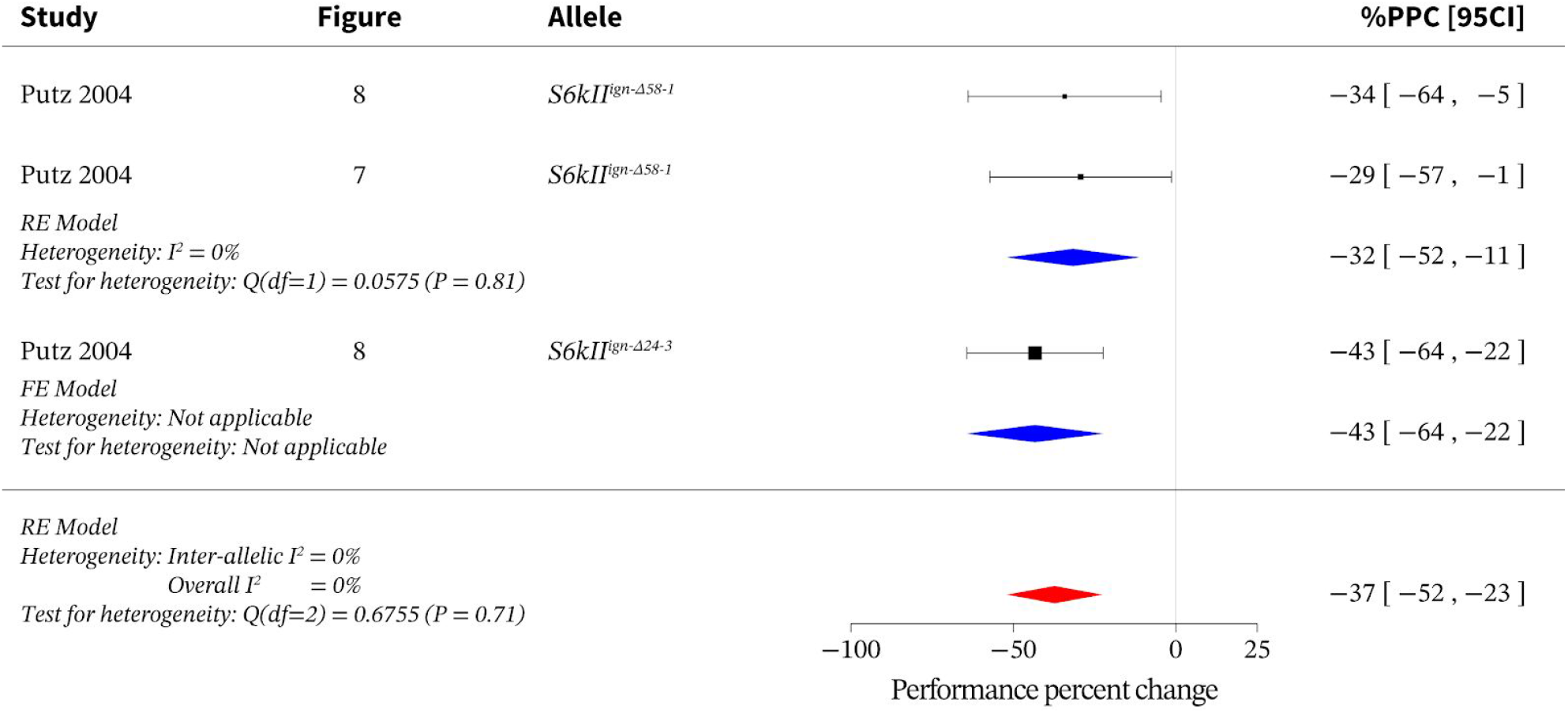
Meta-analysis of loss of function *S6kll* alleles. Meta-analysis of *S6kII* loss-of-function data indicates an overall effect size of –37% [95CI −52, −23] with an overall I^2^ of 0%. FE = fixed effects; PPC = performance percent change; RE = random effects. Blue diamonds represent the effect size of the allelic subgroups of *S6kII.* This data presentation format is repeated for all other forest plots.

**S3.**
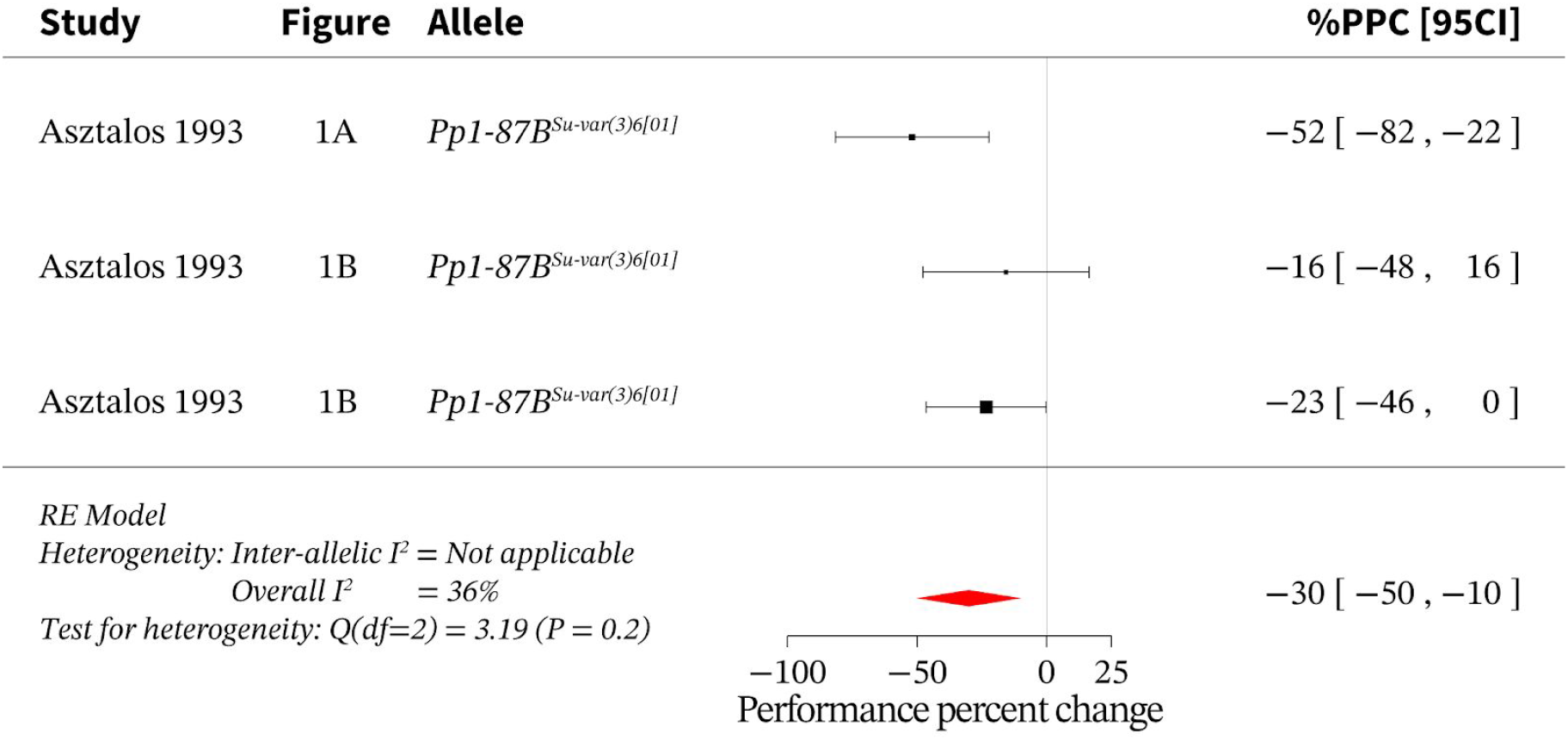
Meta-analysis of loss of function *Pp1-87B* alleles. Meta-analysis of *Pp1-87B* loss-of-function data indicates an overall effect size of –30% [95CI −50, −10] with an overall I^2^ of 36%. PPC = performance percent change; RE = random effects.

**S4.**
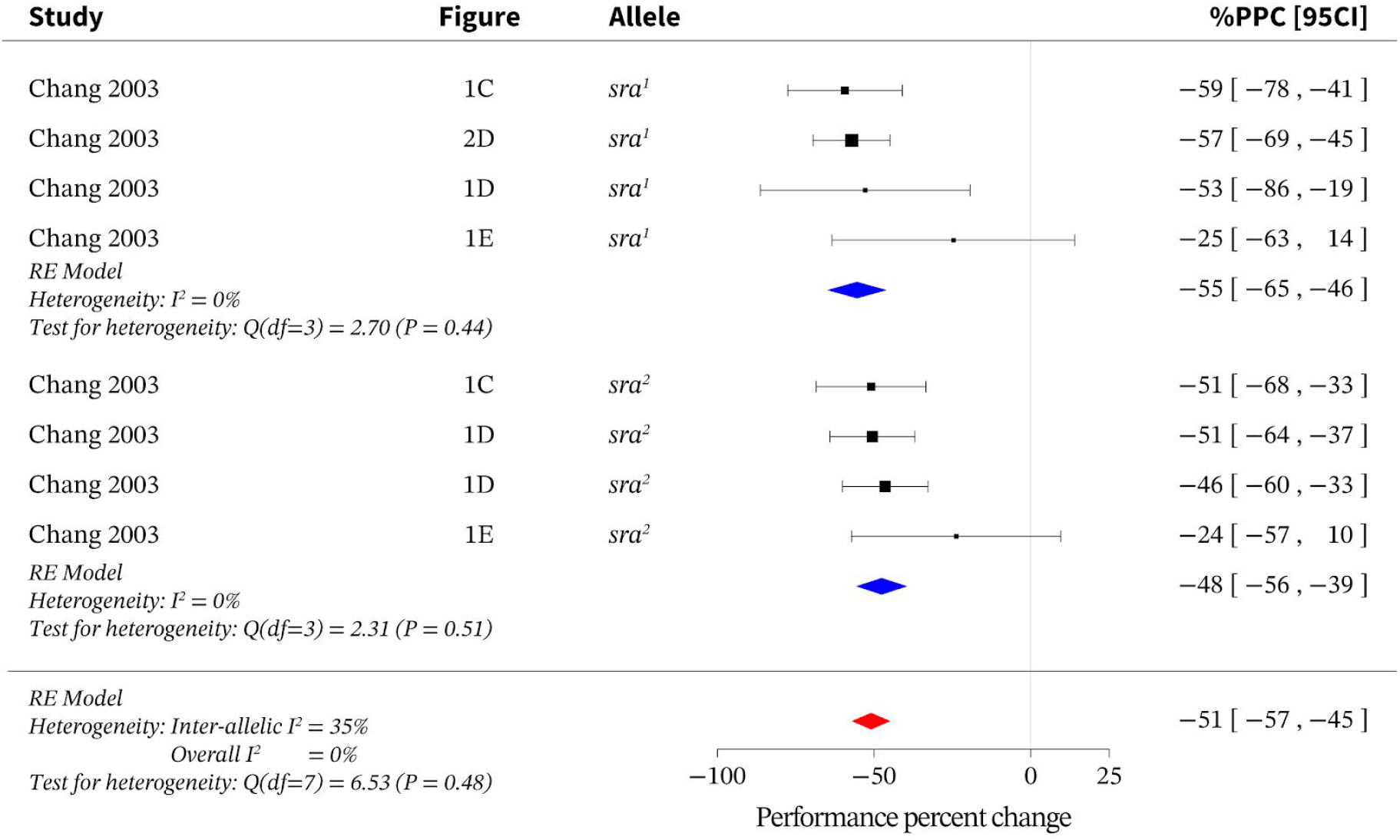
Meta-analysis of loss of function *sra* alleles. Meta-analysis of *sra* loss-of-function data indicates an overall effect size of –51% [95CI −57, −45] with an overall I^2^ of 0%. PPC = performance percent change; RE = random effects.

**S5.**
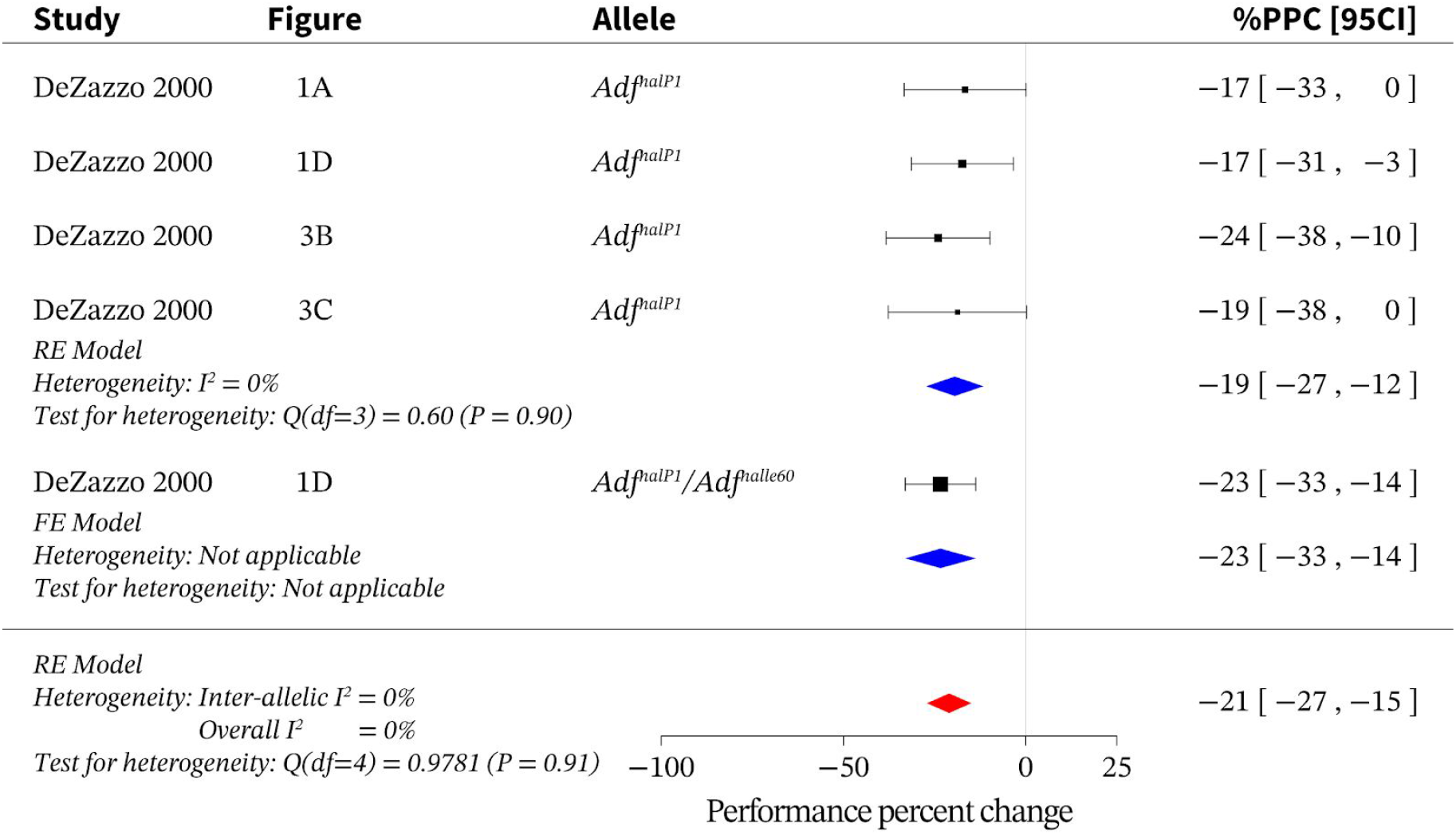
Meta-analysis of loss of function *Adfl* alleles. Meta-analysis of *Adf1* loss-of-function data indicates an overall effect size of –21% [95CI −27, −15] with an overall I^2^ of 0%. FE = fixed effects; PPC = performance percent change; RE = random effects.

**S6.**
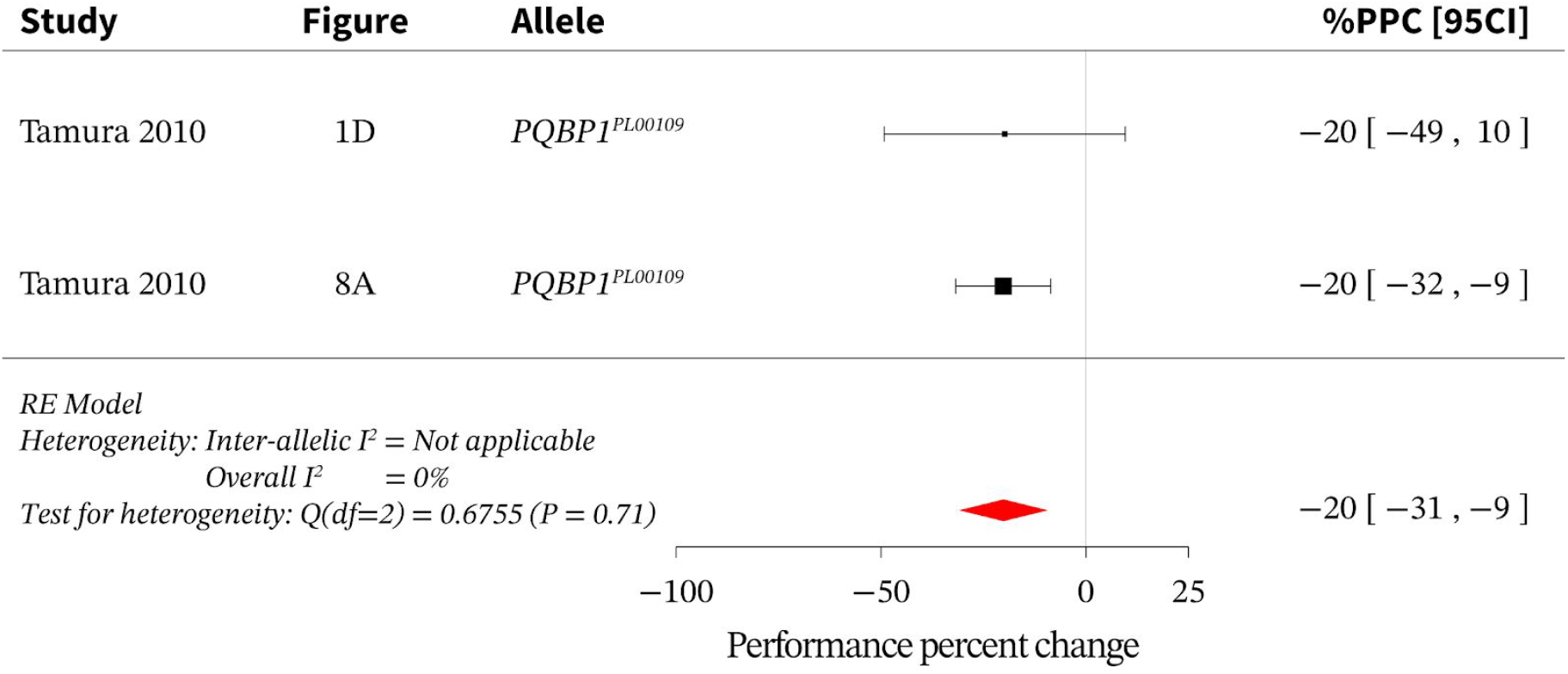
Meta-analysis of loss of function *PQBP1* alleles. Meta-analysis of *PQBP1* loss-of-function data indicates an overall effect size of –20% [95CI −31, −9] with an overall I^2^ of 0%. PPC = performance percent change; RE = random effects.

**S7.**
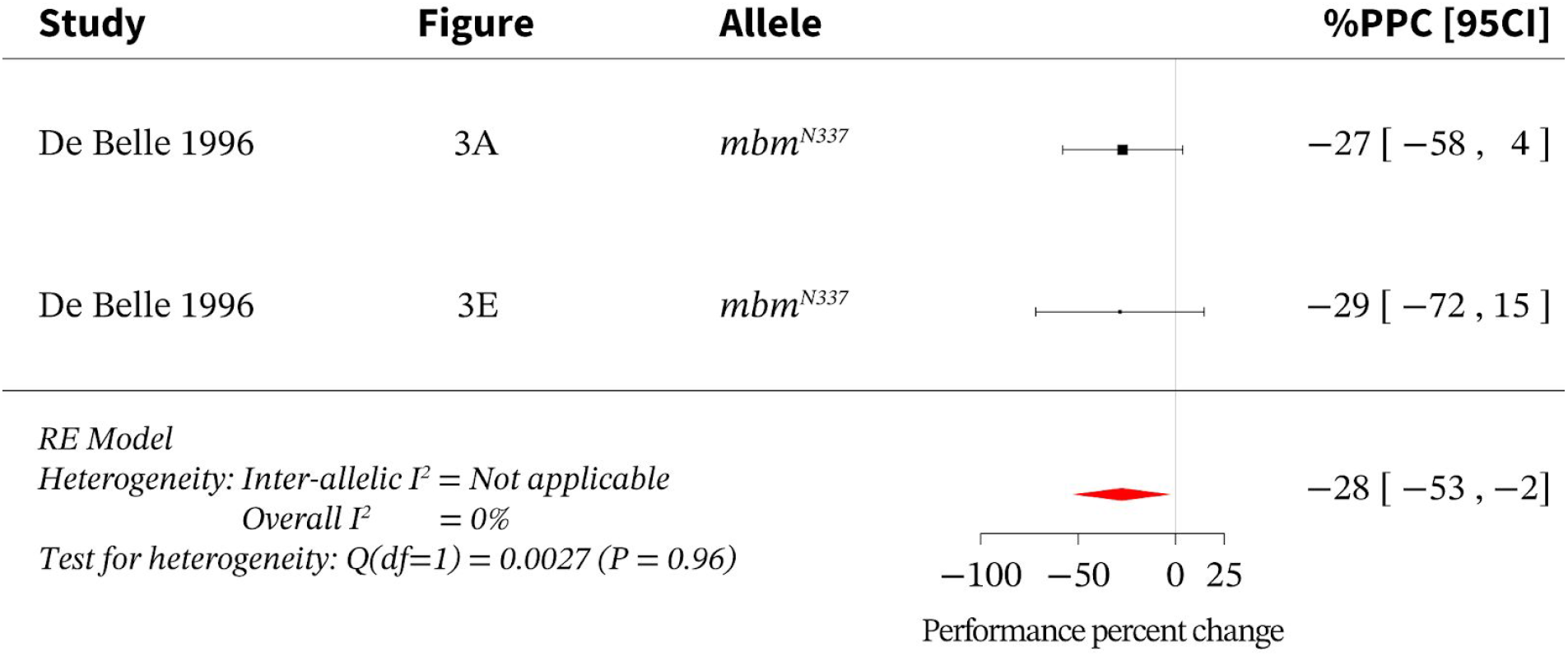
Meta-analysis of loss of function *mbm* alleles. Meta-analysis of *mbm* loss-of-function data indicates an overall effect size of –28% [95CI −53, −2] with an overall I^2^ of 0%. PPC = performance percent change; RE = random effects.

**S8.**
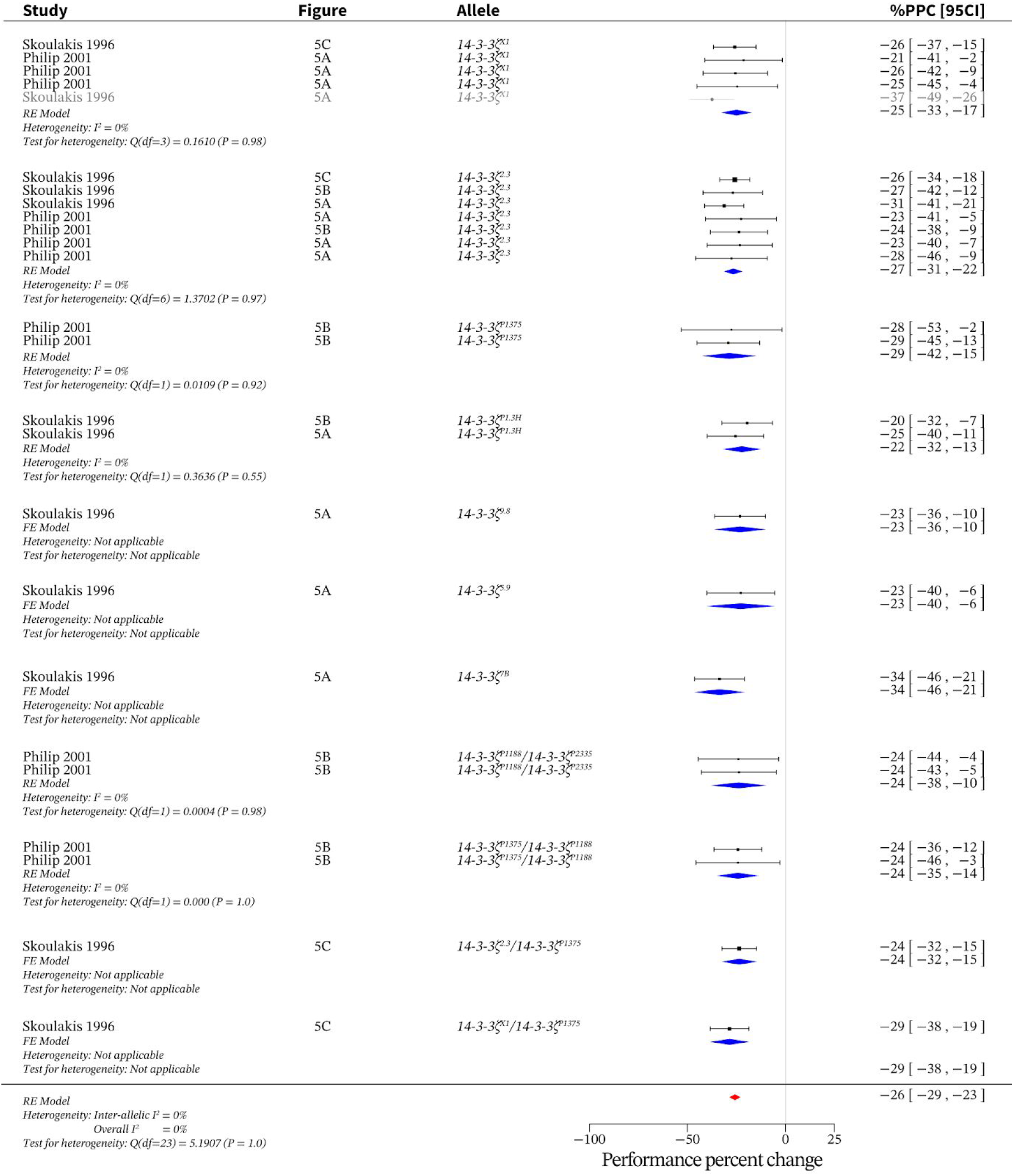
Meta-analysis of loss of function *14-3-3ζ* alleles. Meta-analysis of 74-3-3ζ1oss-of-function data indicates an overall effect size of –26% [95CI −29, −23] with an overall I^2^ of 0%. FE = fixed effects; PPC = performance percent change; RE = random effects. The grey-coloured rows indicate the outliers that were excluded from the calculations based on a Z-score outlier filter (see Methods). This data presentation format is repeated for all other forest plots.

**S9.**
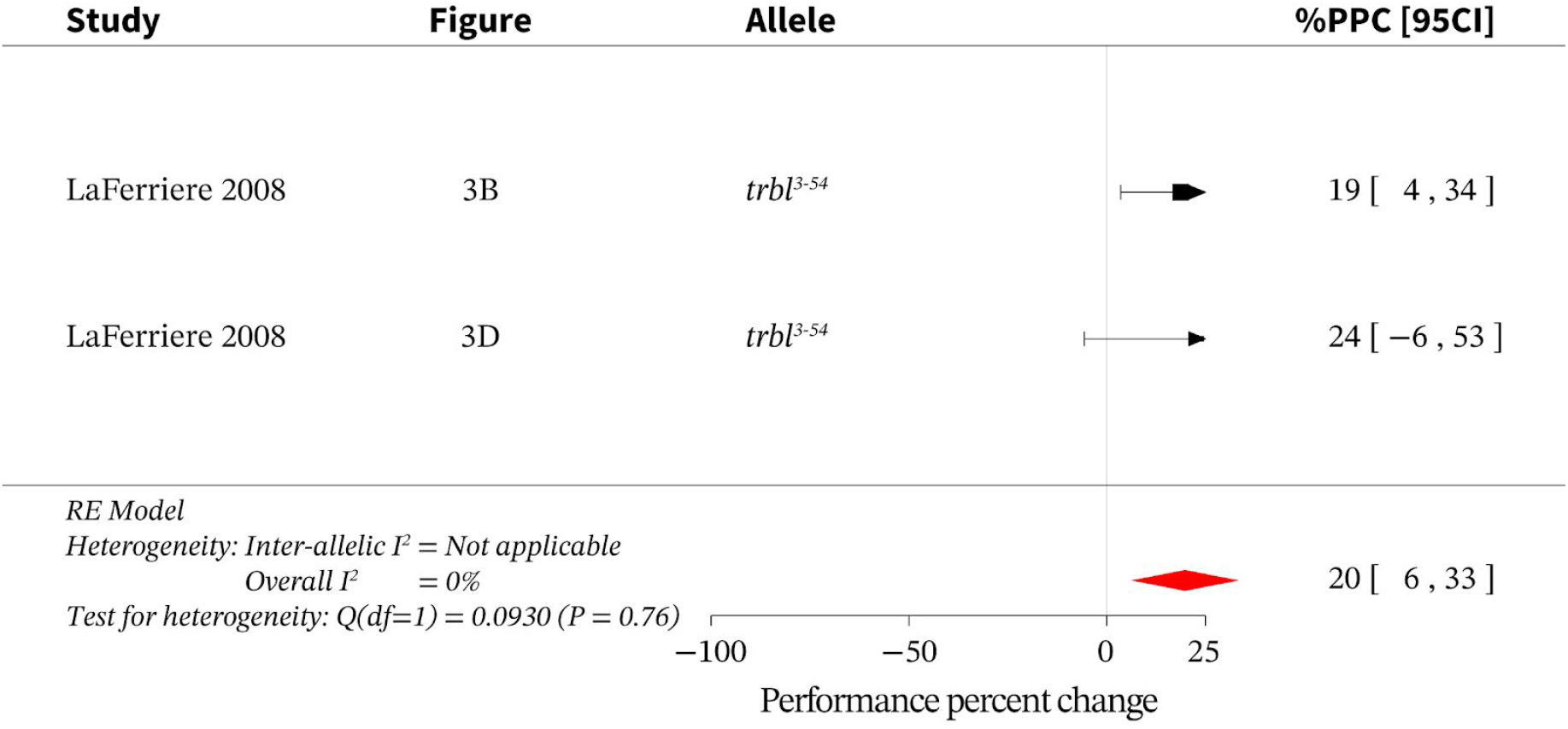
Meta-analysis of loss of function *trbl* alleles. Meta-analysis of *trbl* loss-of-function data indicates an overall effect size of 20% [95CI 6, 33] with an overall I^2^ of 0%. PPC = performance percent change; RE = random effects.

**S10.**
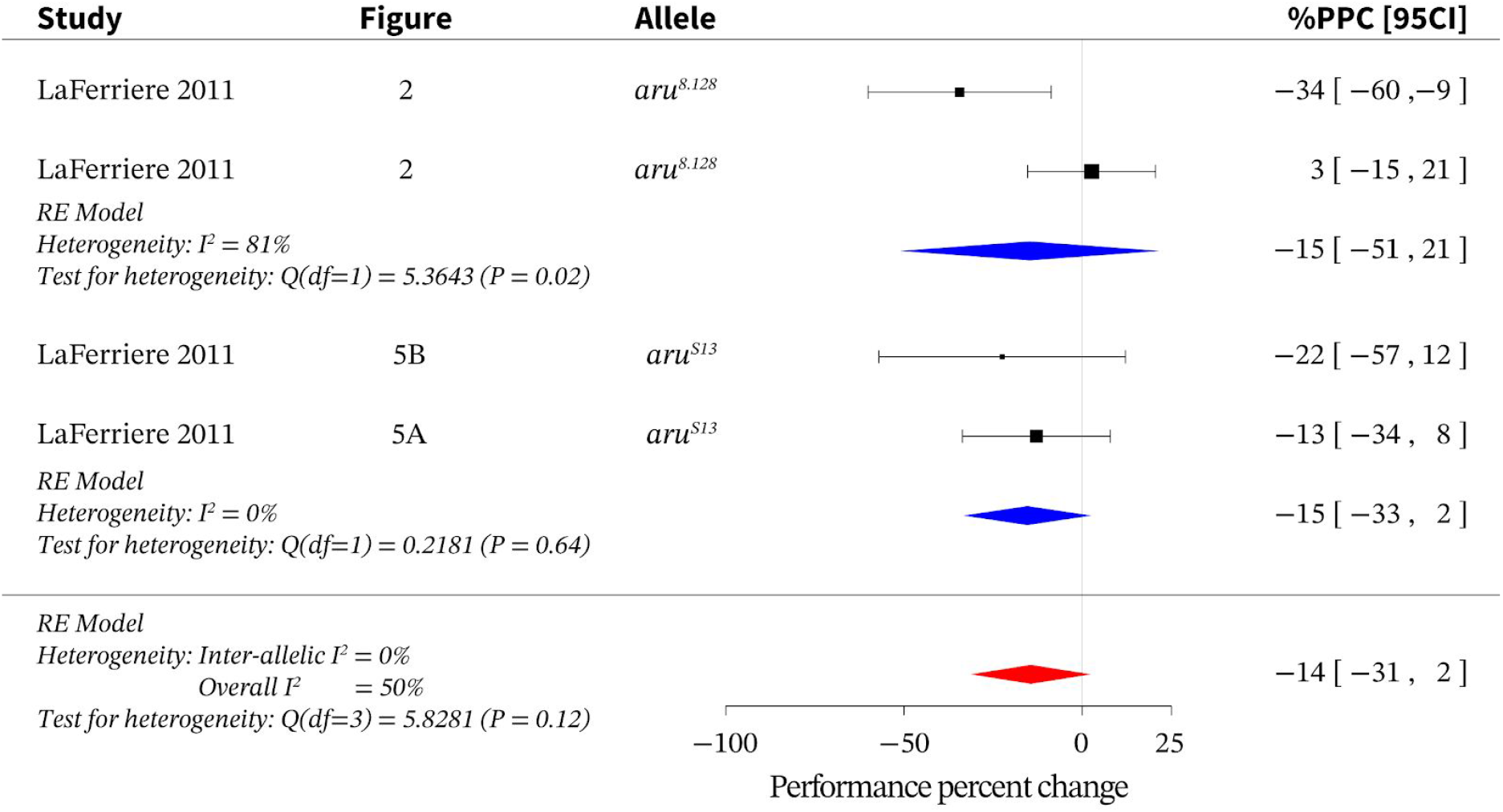
Meta-analysis of loss of function *aru* alleles. Meta-analysis of *aru* loss-of-function data indicates an overall effect size of –14% [95CI-31, 2] with an overall I^2^ of 50%. PPC = performance percent change; RE = random effects.

**S11.**
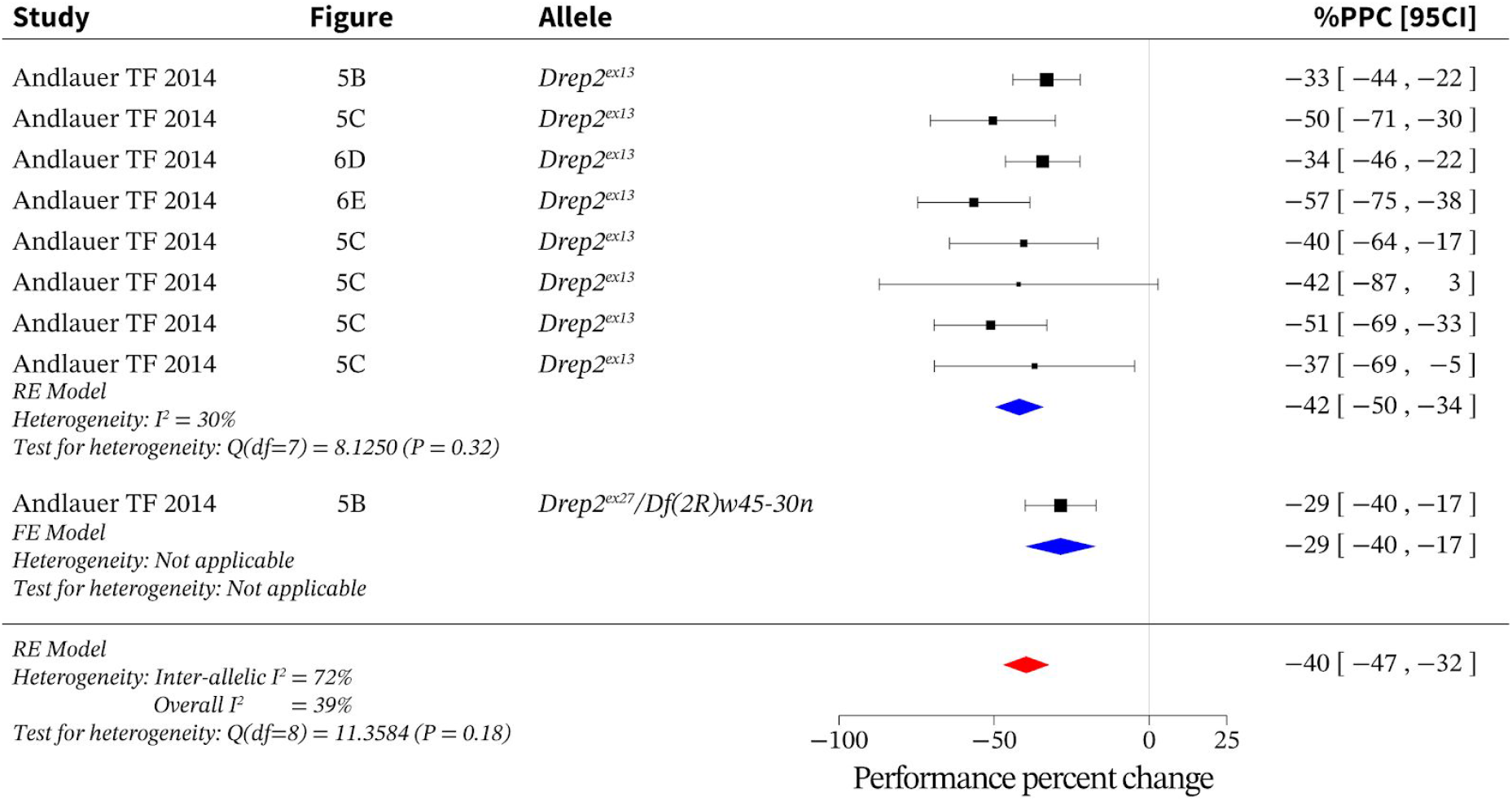
Meta-analysis of loss of function *Drep2* alleles. Meta-analysis of *Drep2* loss-of-function data indicates an overall effect size of –40% [95CI −47, −32] with an overall I^2^ of 39%. FE = fixed effects; PPC = performance percent change; RE = random effects.

**S12.**
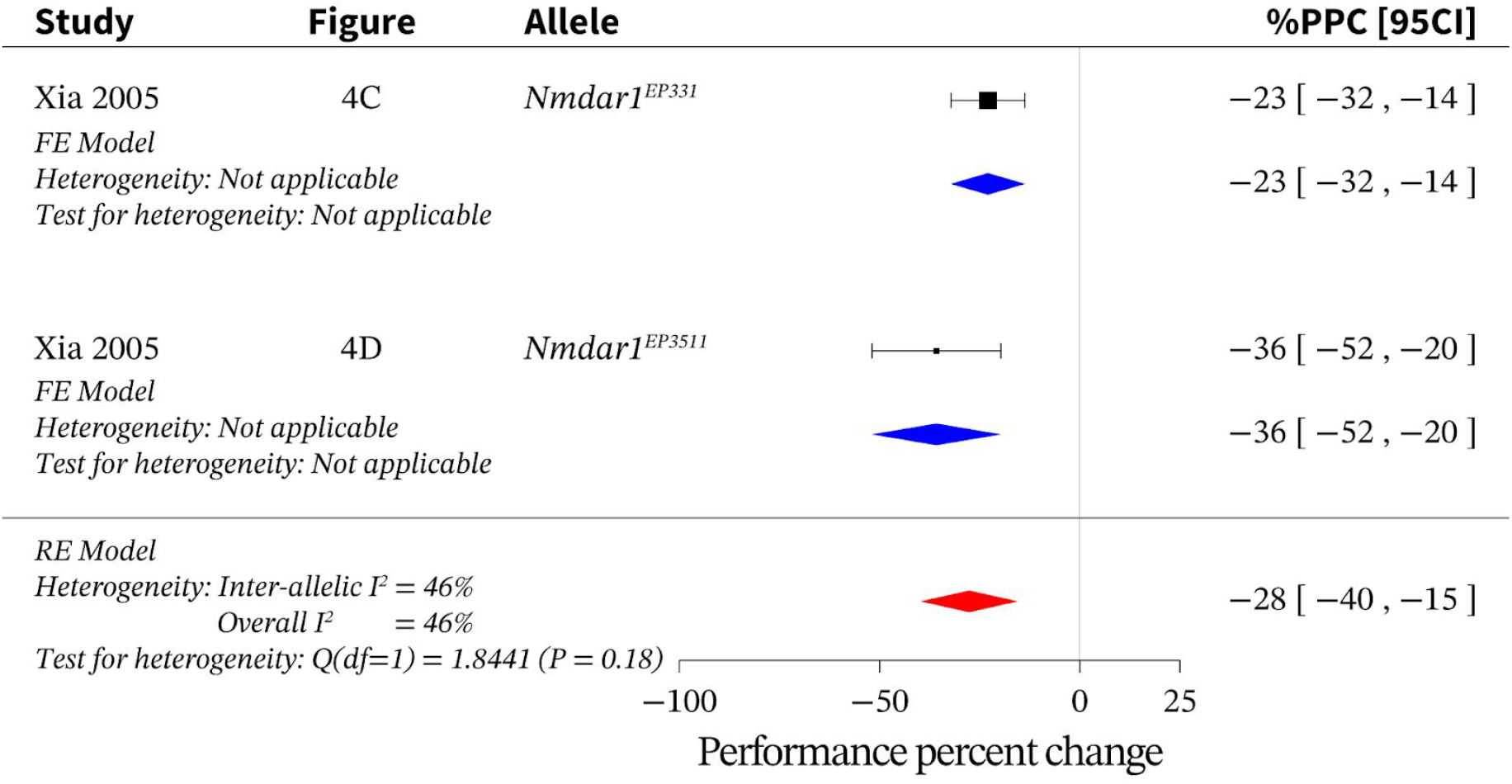
Meta-analysis of loss of function *Nmdar1* alleles. Meta-analysis *of Nmdar1* loss-of-function data indicates an overall effect size of –28% [95CI −40, −15] with an overall I^2^ of 46%. FE = fixed effects; PPC = performance percent change; RE = random effects.

**S13.**
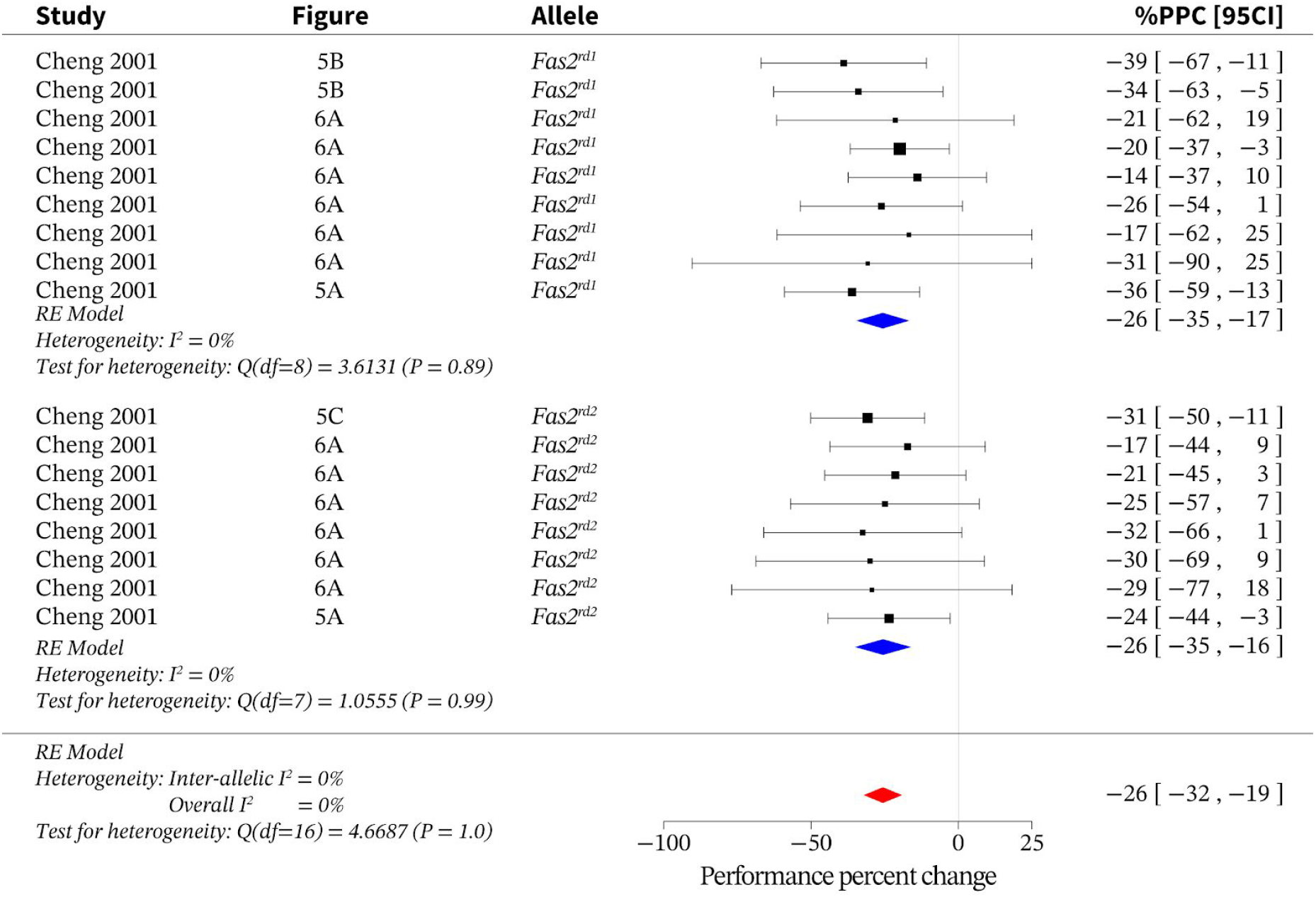
Meta-analysis of loss of function *Fas2* alleles. Meta-analysis of *Fas2* loss-of-function data indicates an overall effect size of –26% [95CI-32, −19] with an overall I^2^ of 0%. PPC = performance percent change; RE = random effects.

**S14.**
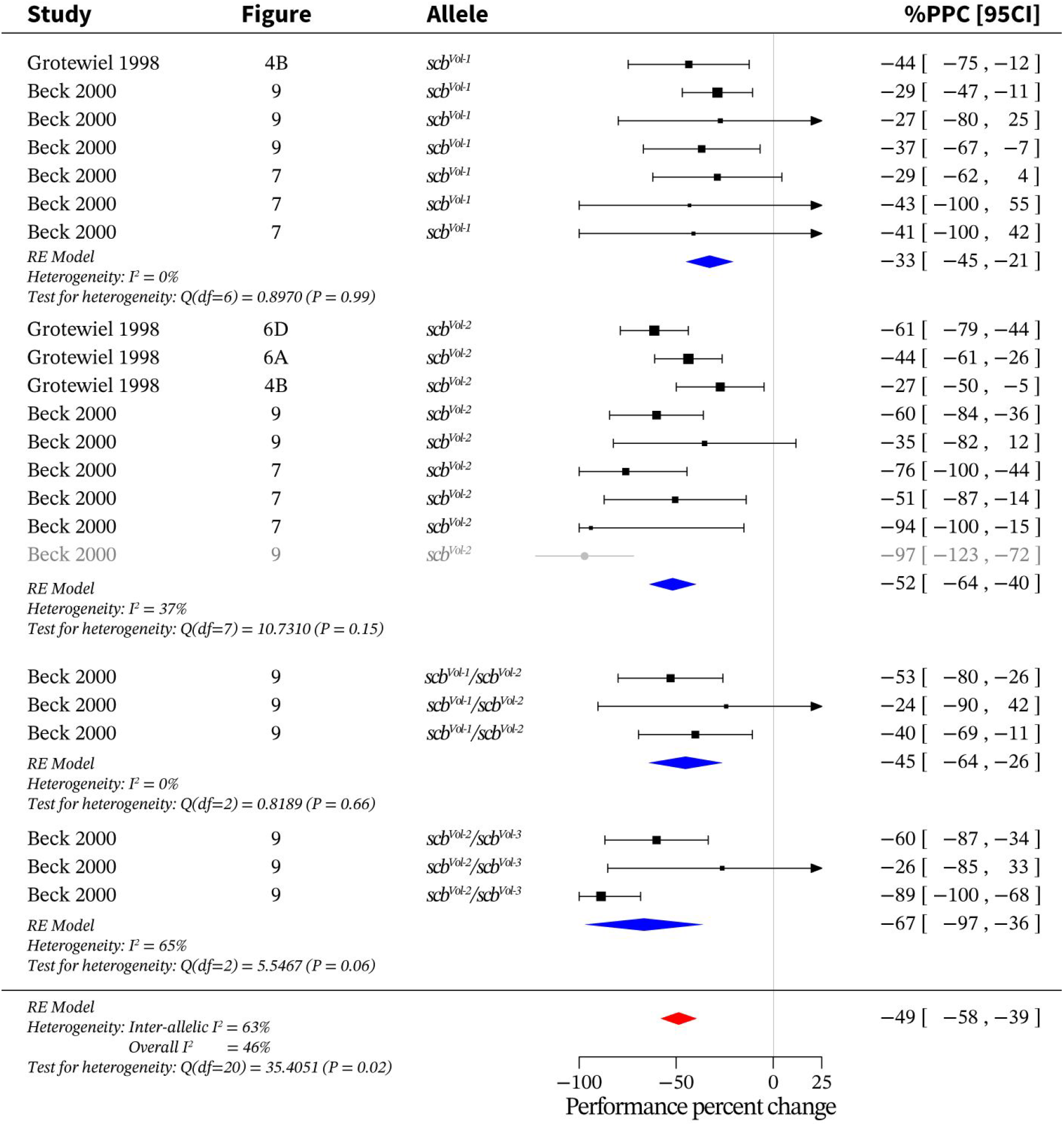
Meta-analysis of loss of function *scb* alleles. Meta-analysis of *scb* loss-of-function data indicates an overall effect size of –49% [95CI −58, −39] with an overall I^2^ of 46%. PPC = performance percent change; RE = random effects.

**S15.**
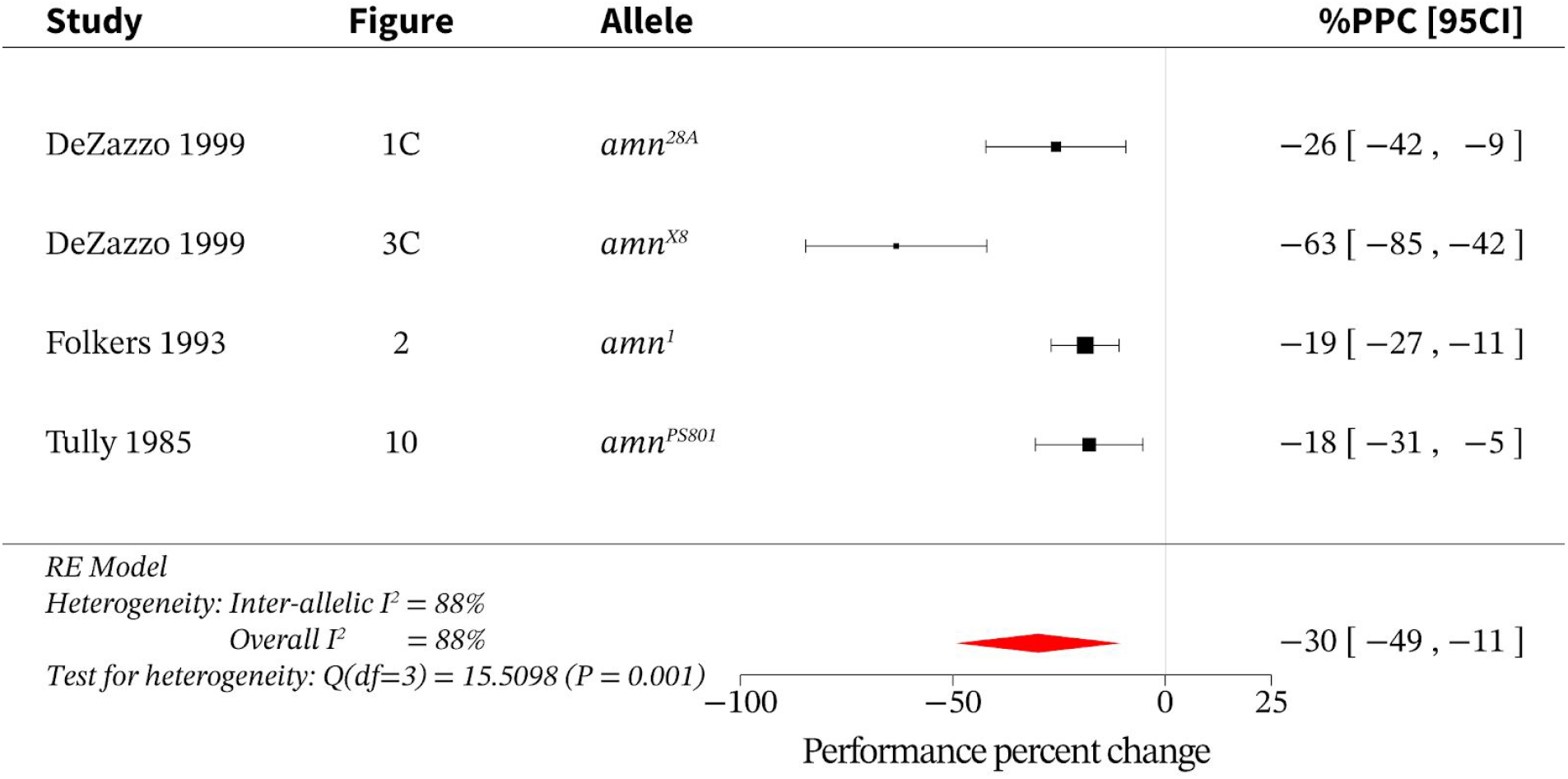
Meta-analysis of loss of function *amn* alleles. Meta-analysis of *amn* loss-of-function data indicates an overall effect size of –30% [95CI −49, −11] with an overall I^2^ of 88%. PPC = performance percent change; RE = random effects.

**S16.**
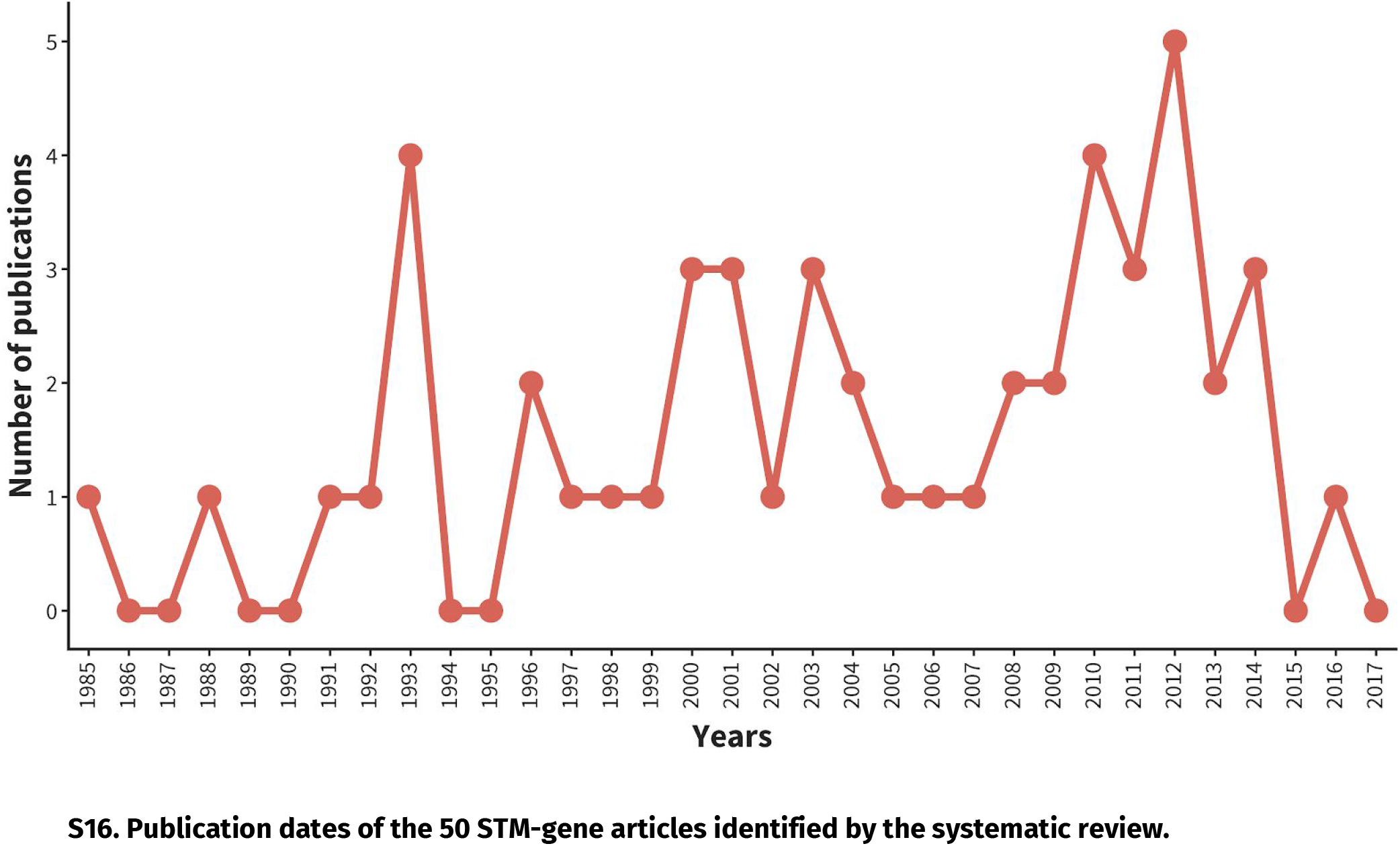
Publication dates of the 50 STM-gene articles identified by the systematic review.

